# Anterior hippocampal integration tracks the developmental emergence of flexible navigation through changing environments

**DOI:** 10.64898/2025.12.11.693744

**Authors:** Nicole L. Varga, Owen W. Friend, Anthony M. Dutcher, Hannah E. Roome, Margaret L. Schlichting, Katherine R. Sherrill, Christine A. Coughlin, Alison R. Preston

**Affiliations:** Department of Neuroscience, The University of Texas at Austin, Austin, TX 78712; Department of Psychology, The University of Texas at Austin, Austin, TX 78712; Senseye, Inc., Austin, TX 78701; Center for Learning & Memory, The University of Texas at Austin, Austin, TX 78712; UX Division, Meta, London, United Kingdom, N1C 4BB; Department of Psychology, University of Toronto, Toronto, ON M5S 3G3; CX Division, Baylor Scott & White Health, Dallas, TX 75240; Department of Psychology, University of Illinois Chicago, Chicago, IL, 60607-7145

**Author notes:** Corresponding Author: Nicole L. Varga Department of Neuroscience; The University of Texas at Austin, 100 East 24th Street; Austin, TX 78712. This manuscript was deposited as a preprint on bioRxiv under an Attribution CC-BY-NC licensing option. Indicates equal contributions (Nicole L. Varga, Owen W. Friend).

**Keywords:** Representational drift, Episodic memory, Development, Inference

## Abstract

Flexible memory depends on cognitive maps that integrate spatial relationships and guide behavior as environments change. The anterior hippocampus is well positioned to support integration across broad spatiotemporal scales, but its late maturation may constrain development of flexible, map-based navigation. Here, we tested whether hippocampal temporal autocorrelation, an index of neural activity stability over time, tracks the development of spatial integration in humans. In a large resting-state fMRI sample (N = 382; ages 5-34 years; 174 females, 208 males), temporal autocorrelation increased with age in anterior, but not posterior, hippocampus. This anterior-specific pattern was replicated in an independent task-based fMRI sample (N = 83; aged 6-12 years and adults; 41 females, 42 males), wherein we linked hippocampal autocorrelation to dissociable components of spatial behavior. The navigation task separated memory for object locations from the ability to update and generalize knowledge across rotations of the distal reference frame and to new objects. Although all age groups learned object locations, only older participants showed evidence that prior spatial structure supported performance as the environment changed across runs. Critically, hippocampal autocorrelation related to behavior only when spatial knowledge was used across runs, rather than improved through within-run feedback, with the clearest profile emerging in adults. In adults, anterior and posterior autocorrelation jointly predicted precise object-location memory, whereas anterior autocorrelation uniquely predicted efficient navigation trajectories. These findings identify anterior hippocampal temporal autocorrelation as a later-maturing computation that supports the transition from local spatial learning in childhood to flexible navigation through changing environments in adulthood.

**Significance Statement:** Finding our way through the world requires more than remembering where things are. We also need to use what we have learned to take new routes, adjust when familiar places change, and apply old knowledge to new situations. These abilities improve from childhood to adulthood, but the brain changes that support this transition remain unclear. We show that a signal in anterior hippocampus, a brain region important for linking experiences, becomes more stable over development. Using a navigation task that separated remembering object locations from flexibly using a map, we found that this signal was most strongly tied to adults’ ability to navigate efficiently through changing environments. These findings reveal a hippocampal mechanism that supports flexible navigation as individuals mature.

## Introduction

Memory enables recall of individual events as well as their integration into cognitive maps—representations that encode holistic relationships among locations (Tolman, 1948), allowing individuals to infer novel routes and generalize knowledge across contexts (Behrens et al., 2018). Anterior hippocampus, with its broad spatial fields (Jung et al., 1994) and extended temporal integration window (Kjelstrup et al., 2008; Royer et al., 2010), is well-suited for such representations (Strange et al., 2014). However, because this region matures relatively late (Demaster et al., 2014, 2016; Vijayarajah & Schlichting, 2026), children and adolescents may be less able to form and generalize cognitive maps. Although hippocampal maturation has been linked to improved memory (Ghetti & Bunge, 2012) and inference (Schlichting et al., 2016), how anterior hippocampal development supports map-based navigation remains unknown.

Behavioral evidence suggests a developmental shift from representing individual experiences to integrating information across time and space (Bauer et al., 2021; Shing et al., 2019; Varga et al., 2019). Children retrace directly learned routes, whereas adolescents infer novel shortcuts (Burles et al., 2020) and integrate separate paths (Nazareth et al., 2018), with continued refinement into adulthood (Brucato et al., 2022). This increased flexibility parallels developmental change in anterior hippocampus structure (Schlichting et al., 2016), activation (Demaster et al., 2016), and connectivity (Calabro et al., 2020) that is more protracted than that of posterior hippocampus (Xie et al., 2024) and may promote mature memory integration (Schlichting et al., 2015; Varga et al., 2024). Here, we test how spatiotemporal integration along the hippocampal long axis changes with age and enables flexible navigation.

A leading view (Raut et al., 2020) proposes that spatiotemporal integration is supported by temporal autocorrelation (Paz et al., 2010)—the stability of neural activity patterns over time, which may support linking of temporally distinct events into integrated representations (Varga & Manns, 2021). Adults show greater temporal autocorrelation in anterior than posterior hippocampus during navigation (Brunec et al., 2018). Moreover, greater hippocampal autocorrelation during learning predicts stronger clustering of related events in memory (Rait et al., 2025), whereas disruptions in hippocampal autocorrelation reflect a shift away from maintaining an existing representation toward encoding new information (Sinclair et al., 2021). We thus predicted that anterior hippocampal autocorrelation would increase with age, more than posterior hippocampus, and would further promote navigation by enabling individuals to deploy an existing cognitive map to navigate a changing environment.

First, we leveraged a large resting-state fMRI sample to examine developmental differences in hippocampal temporal autocorrelation from childhood through adulthood. Although prior studies have shown that children and adolescents recruit anterior hippocampus less than adults during memory encoding (Ghetti et al., 2010) and retrieval (Demaster et al., 2016), these investigations have largely focused on activation magnitude rather than underlying representations (Benear et al., 2022; Schlichting et al., 2022; Varga et al., 2025). Moreover, prior work has characterized developmental differences in hippocampal representations using measures of their spatial organization (Callaghan et al., 2021). However, it remains unknown whether development is also associated with changes in the temporal stability of hippocampal representations, as indexed by temporal autocorrelation. Such changes may reflect maturation of integrative computations that support broader and more flexible memory representations.

With a second, independent study, we examined how hippocampal integration impacted cognitive map formation and expression. Specifically, we asked whether greater anterior hippocampal autocorrelation is associated with a shift from behaviors anchored to individual object-location associations, thought to rely more on posterior hippocampus (Evensmoen et al., 2013, 2015), toward flexible, map-based navigation that represents the relationships among locations and environmental structure (Javadi et al., 2017). We tested whether greater anterior autocorrelation is associated with more efficient navigation through a changing—but structurally intact—environment, reflecting maintenance and expression of an integrated cognitive map. By linking developmental differences in the stability of hippocampal representations to distinct navigation behaviors, the present work identifies a key neural mechanism supporting the transition from representing isolated experiences to constructing integrated, map-like knowledge.

## Materials and Methods

### Participants

#### Resting-state sample

In the resting-state study, we pooled a large sample of child, adolescent, and adult data to investigate the development of hippocampal temporal autocorrelation. Prior work shows that resting-state temporal autocorrelation is stable across repeated within-person assessments in adults, suggesting that task-independent autocorrelation reflects an intrinsic signature of individual brain function (Bouffard et al., 2023). As such, it may provide a useful window into how hypothesized integrative computations change with age.

To assemble a large, continuous developmental sample, we aggregated data from three existing resting-state datasets: the open-source Healthy Brain Network (HBN; Releases 1-6; (Alexander et al., 2017), the Nathan Kline Institute-Rockland Sample (NKI-RS; Initial Release; (Nooner et al., 2012), and a previously published dataset collected at the University of Texas at Austin (UT Austin; Schlichting et al., 2016).

For the open-source HBN (n = 333) and NKI-RS (n = 90) datasets, participants were included if they were between 5 and 34 years of age and had previously passed data quality validation (Cohen et al., 2022). From this subset, 87 additional participants (84 HBN; 3 NKI-RS) were excluded due to excessive head motion based on the stricter thresholds applied in the present study (see below; **MR image preprocessing**). The remaining 336 (249 HB; 87 NKI-RS) participants (5.66 - 34.0 years; 151 females, 185 males) were included in the final resting-state sample. Additional demographic details for the HBN and NKI-RS samples are reported elsewhere (Alexander et al., 2017; Nooner et al., 2012).

Imaging data for the UT Austin resting-state sample was collected under protocols approved by the Institutional Review Board at UT Austin. Individuals were recruited from the local community and received monetary compensation for participation. Following informed consent/assent, participants were screened for: (1) basic inclusion criteria, including no psychiatric conditions, right handedness, no color blindness, and native English speakers; (2) neuropsychological conditions using the Child Behavior Checklist for minors (CBCL; Achenbach & Edelbrock, 1991) and the Symptom Checklist 90-Revised for adults (SCL-90-R; Derogatis & Cleary, 1977); and (3) general intelligence using the Wechsler Abbreviated Scale of Intelligence (WASI-II; Wechsler, 2011). Participants were excluded if they did not meet basic inclusion criteria, scored more than one standard deviation (SD) above the normative mean on neuropsychological measures, or scored more than two SDs below the mean on the full-scale IQ composite.

Of the 90 initially recruited UT Austin participants, 20 were excluded prior to the MR portion due to the presence of a psychiatric condition (n = 19 minors, 1 adult), handedness (n = 1 minor), neuropsychological screening criteria (n = 2 minors, 9 adults), or withdrawal (n = 3 minors, 4 adults). In addition, 24 participants were excluded due to data quality issues, including poor quality anatomical data (n = 1 minor; 2 adults) and excessive head motion (n = 21 minors; see **MR image preprocessing** below). The remaining 46 participants (6.92 - 28.83 years; 23 females, 23 males) were included in the final sample.

Following exclusions, the final resting-state sample consisted of 382 individuals (*M* Age = 13.69 years, *SD* = 5.18 years, *Range* = 5.7-34.0 years; 174 females, 208 males). The sample size is consistent with past developmental work using resting-state functional MRI to examine age-related differences along the hippocampal long axis (Xie et al., 2024). Also consistent with prior developmental imaging work, head-motion-related exclusions were more common at younger ages, potentially resulting in a final developmental sample that overrepresents individuals who were better able to remain still during scanning. Age distributions by sample are provided in Supplemental Figure 1 (**Figure S1, Panel A**).

#### Task-based navigation sample

To test the functional significance of developmental differences in anterior hippocampal integration, we next examined whether a more mature pattern of anterior hippocampal autocorrelation was associated with more sophisticated navigation behavior. Importantly, the task-based navigation sample also provided a test for replication of the developmental pattern observed at rest. Specifically, because spatial navigation preferentially engages anterior hippocampal autocorrelation relative to posterior hippocampal autocorrelation in adults (Brunec et al., 2018), this task allowed us to test whether the predicted developmental pattern of lower anterior temporal autocorrelation was evident at rest as well as in a behavioral context that strongly engages this anterior hippocampal computation.

We tested this question in an independent sample of participants recruited from the local UT Austin community who were screened using the same criteria described in the UT Austin resting-state sample above. For this study, participants were recruited within three approximately equal age groups comprising children (6-9 years), early adolescents (10-12 years), and adults (18-34 years). This sampling strategy was motivated by prior behavioral studies that have identified a shift toward more adult-like cognitive map formation around 10-12 years of age (Brucato et al., 2022; Burles et al., 2020; Nazareth et al., 2018), allowing us to test for a corresponding shift in hippocampal autocorrelation before, during, and after this behavioral transition.

Our target sample size for each age group (*N* = 25) was guided by prior, related research in adults (Brunec et al., 2018), which showed that this number was sufficient to detect differences in temporal autocorrelation between the anterior and posterior hippocampus during spatial navigation. The sample size also aligns with past developmental work that has demonstrated age-related differences in hippocampal representation using multivariate methods (Callaghan et al., 2021; Varga et al., 2025).

Of the 145 participants initially recruited, 62 individuals were excluded due to a psychiatric condition (n = 1 minor), non-native English status (n = 2 minors), neuropsychological screening (n = 8 minors), voluntary withdrawal (n = 18 minors, 1 adult), technical difficulties during image acquisition (n = 6 minors), failure to complete a sufficient number of runs (n = 11 minors, 1 adult; Minimum = 2 runs), or excessive head motion (n = 14 minors). The remaining 83 participants (*M* Age = 13.8 years, *SD* = 5.9 years, *Range* = 6.9-33.4 years; 41 females, 42 males; children: 6-9 years, *N*=26; early adolescents: 10-12 years, *N*=29; adults: 18-34 years, *N*=28) were included in the final task-based sample.

As in the resting-state sample, younger participants were more likely to be excluded for head motion or additional reasons, such as incomplete task data. Importantly, while this potential selection bias should be considered when interpreting the findings, it is noteworthy that its likely effect would be to attenuate, rather than inflate, any developmental differences between younger participants and adults. That is, because the retained younger participants were potentially more similar to adults, at least in their ability to meet the scanning and experimental task demands, this selection would make developmental differences more difficult—not easier—to detect. The full age distribution is provided in Supplemental Figure 1 (**Figure S1, Panel B**).

### Procedures for resting-state study

All resting-state scans were collected within study sessions that included several additional scans and/or tasks. For resting-state scans, participants were instructed to lie in the scanner and stay as still as possible. Participants at the HBN and UT Austin sites provided two runs of resting-state data each lasting five minutes, while participants at the NKI-Rockland site provided one ten-minute run of resting-state data (see below; **MR image acquisition**).

### Stimuli and materials for navigation study

The spatial memory paradigm was adapted from prior adult research (Doeller et al., 2008) for use with children and adolescents. A custom 3D virtual environment (**Figure 1**) was built in Unity (version 2017.5.2.2; Unity Technologies, 2017). The environment consisted of a circular meadow arena (67.3 virtual meters [vm] in diameter) enclosed by a 3-vm-high wall with invisible boundaries. Four distal cues (mountains) were positioned at the cardinal directions surrounding the arena, and a proximal landmark (traffic cone) was positioned inside the arena 45° between two mountains.

**Figure 1.**
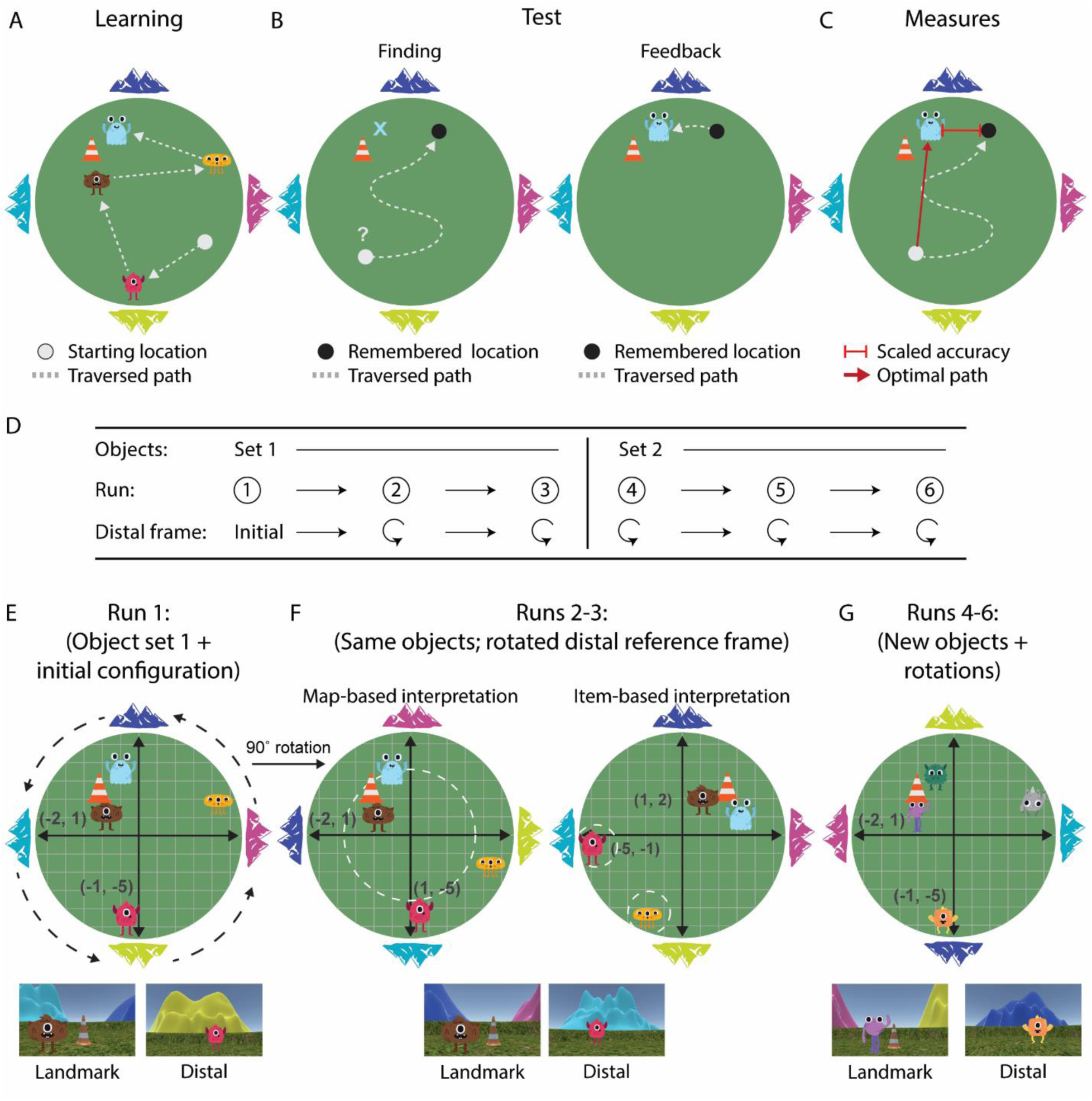
Schematic of navigation task and manipulations. **A**) During learning trials, participants were spawned into the virtual environment and could see the first object at its location. They were instructed to move toward that visible object to learn its precise positioning within the environment. Once they navigated to the visible object, it disappeared and the second object became visible. They then moved toward it, following this procedure until they learned the locations of each object. **B**) During Test trials, participants were spawned into one of eight starting positions and were cued to find one of the target objects (not currently visible to them; i.e., Finding). Once in the remembered location, the participants pressed a button and then the object appeared. Participants then navigated to the target object’s visible location (i.e., Feedback). **C**) For each trial, we measured the accuracy of the participant’s location memory, operationalized as scaled accuracy—i.e., the distance error in virtual meters (vm) between the remembered location and the object’s true coordinate. We also measured excess path, calculated as the difference between a maximally efficient Euclidean path between the spawn and target locations and the total distance (finding + feedback) traversed on that trial. **D)** Summary of the across-run environmental manipulations. The table indicates the transition from Object Set 1 to Object Set 2 (between Runs 3 and 4), as well as the rotation of the distal reference frame across Runs 2-6. **E**) The initial object configuration was **F)** systematically rotated either 90 or 180 degrees across runs, preserving the spatial relationship among objects, but resulting in new distal associations for each object. If participants formed holistic representations of the environment (left arena, map-based interpretation), they could recognize that the relationship among objects as well as each object’s relationship with the proximal cue remained constant; preserving the initial coordinates. If participants did not detect this common spatial structure (right arena, item-based interpretation), they would relearn new local coordinates for each individual object **G**) Object sets changed between runs 1-3 and 4-6 while the spatial configuration and across-run rotations remained. Thus, participants could generalize their internal spatial map to the new object set, thereby reducing the demands of learning new local coordinates on each run across the task.

Eleven object cues were used across the experiment (**Figure 1A; 1G**). Each object consisted of a multicolored, unfamiliar cartoon character. Eight were used in the main spatial memory task (four in object set 1 and four in object set 2; see below), one object was used in navigation control trials, and two objects were used during practice.

### Procedures for task-based navigation paradigm

While the resting-state study sought to establish whether anterior hippocampal temporal autocorrelation increases from childhood to adulthood, the task-based navigation study examined the behavioral implications of developing temporal autocorrelation patterns. Specifically, we tested the hypothesis that developmental increases in anterior hippocampal autocorrelation would support integration across increasingly broad spatiotemporal scales, enabling the formation of holistic spatial representations—or cognitive maps—that link experiences across space and time (Behrens et al., 2018; Brucato et al., 2022; Burles et al., 2020; Nazareth et al., 2018; Tolman, 1948). Such representations, in turn, should facilitate flexible navigation and generalization across changing spatial contexts (Behrens et al., 2018; Burles et al., 2020; Tolman, 1948).

To test these predications, we used the virtual environment and object sets described above to implement an adapted spatial memory task (Bullens et al., 2010; Doeller et al., 2008), in which starting locations and environmental structure were systematically manipulated across trials and runs. Importantly, temporal autocorrelation has been shown to be modulated by task demands in adults (Brunec et al., 2018), with greater anterior hippocampal temporal autocorrelation observed during novel, more flexible navigation conditions. This design thus allowed us to assess developmental differences in flexible navigation behavior and test whether these differences were related to developmental variability in anterior hippocampal autocorrelation, in particular.

#### Task structure

Participants completed six runs of the task. In each run, they navigated the environment to locate four target objects (“friendly monsters”) arranged relative to environmental cues (**Figure 1A**). Two objects were positioned near the proximal landmark (the traffic cone), with one located closer to the landmark and one farther away. The remaining two objects were positioned near the distal cues (the mountains), with one centered in front of a mountain and one offset to the side of a mountain.

Each run consisted of learning (**Figure 1A**) and test (**Figure 1B**) trials, with test trials comprising finding and feedback phases. During learning trials (4 per run; **Figure 1A**), participants were spawned at a starting location within the environment and could see the first object at its location. Participants were instructed to move toward that visible object to learn its precise positioning within the environment. Once they navigated to the object, it disappeared and the second object became visible. They then moved toward that object, following this procedure until they learned the locations of each of the four objects.

During the subsequent test trials (12 per run; **Figure 1B**), participants were instructed to find the target objects (not currently visible to them). Each test trial began with presentation of the target object on a gray screen, followed by a jittered fixation (2-6s; mean = 4s). Participants were then placed at one of eight pseudo-randomized starting locations in the environment (**Figure S2, Panel A**) and navigated to the remembered location (“Finding” phase: **Figure 1B, left arena**), pressing a button when they had reached it (max search time = 60s). Following this response, the true object location was revealed, and participants navigated to it as corrective feedback (“Feedback” phase; **Figure 1B; right arena**). The 12 test trials in each run were organized into three blocks of four trials, with each block including one trial per object in randomized order. Thus, within a run, participants were cued to find each object three times, each from different starting positions.

For each test trial, we assessed two complementary measures of spatial knowledge (**Figure 1C**). First, we measured the precision of a participant’s object-location memory, quantified as distance error—i.e., the distance (in vm) between the remembered location and the object’s true location. These values were transformed into a scaled accuracy metric to account for variation in chance-level performance across locations in the circular environment (**Supplemental Materials: Scaled accuracy calculation**; **Figure S2, Panel B**; see also Bullens et al., 2010), with higher values reflecting greater accuracy.

Second, we measured the efficiency of a participant’s trajectory from the pseudo-random starting position to the target location, referred to as excess path, or the difference between the shortest Euclidean path and the total path length traversed by the participant (including finding and feedback; **Figure 1C**; **Supplemental Materials: Excess path calculation**). Lower values for excess path reflect more efficient navigation trajectories through the environment. Importantly, these measures dissociate two potential navigation strategies: (1) a local, item-based approach, in which each object is encoded in relation to nearby environmental cues, and (2) a global, cognitive map-based approach, in which spatial relationships among environmental cues and objects are integrated into a holistic representation of the environment. Scaled accuracy (derived from distance error) could be supported by either posterior hippocampal coding of precise object locations and/or anterior hippocampal integration of each object within the broader spatial structure. By contrast, excess path provides a more stringent index of anterior hippocampal contributions to map-based navigation because efficient trajectories from novel starting positions require participants to locate themselves within the global environment and plan direct routes through that space.

Participants navigated from a first-person perspective (1.7 vm height) at a constant speed (8 vm/second) using a button box with keys for left, forward, and right movement. Prior to entering the scanner, participants completed extensive practice with the navigation procedure on a separate laptop in a separate square desert arena that was comparable in size to the main task arena but contained distinct environmental cues. Specifically, the practice arena was surrounded by four distal mountains that differed in formation and color from those in the main task and contained a rotationally symmetrical trash can as a proximal landmark cue. Participants were first walked through the different components of the task and practiced navigating to two monsters (target objects), one associated with the proximal landmark and the other with a distal cue. This practice phase provided participants with an opportunity to learn and demonstrate comprehension of the task procedures before entering the scanner.

Once in the scanner, each trial block was preceded by a control trial designed to assess participants’ basic ability to navigate using the button box. During these trials, participants navigated toward a visible object within a separate open-field environment that contained no environmental cues and was rendered to infinity. The control trials therefore provided an in-scanner assessment of basic mastery of the motor controls, while also providing an ongoing measure of engagement throughout the experiment. This was particularly important for younger participants, who might take longer to retrieve a target location during the main task and therefore spend less time actively navigating, despite remaining engaged with the task. Moreover, because navigation performance could vary across trial blocks as participants strengthened memories for new object-location associations, including a control trial before each block allowed us to monitor navigation ability repeatedly throughout the run, including early on when new object-location memories were being formed. Custom MATLAB code (version 2014b) was used to generate condition files for input to Unity and to record behavioral data.

#### Description of environmental change across runs

Our primary goal was to determine how developmental differences in hippocampal autocorrelation relate to flexible navigation behaviors that depend on cognitive maps. To this end, as discussed above, we incorporated behavioral measures (**Figure 1C**) that dissociate between two potential navigation strategies: (1) a local, item-based approach, in which each object is encoded in relation to nearby environmental cues, and (2) a global, map-based approach, in which spatial relationships among environmental cues and objects are integrated into a holistic representation of the environment. Moreover, we implemented complementary within- and across-run task manipulations designed to test three core properties of cognitive maps—flexible navigation, updating, and generalization.

Together, these manipulations allowed us to distinguish between hippocampal subsystems supporting local versus global spatial learning. Limited integration across time and space—characteristic of an immature hippocampus in which only posterior autocorrelation is fully functional—should lead to a local, item-based approach, resulting in precise but inefficient navigation. In contrast, mature anterior hippocampal integration should enable representation and extension of global environmental structure, allowing accurate and efficient navigation, even following changes in task contingencies. We detail each of these changing task contingencies in turn.

##### Flexible navigation from novel starting positions

A defining property of cognitive maps is the ability to support efficient navigation to known goal locations via novel routes that have not been directly experienced. This ability depends on forming a holistic map of the environment that encodes both the relationships among stable landmarks (e.g., the relative positions of the mountains), as well as the relationship between these environmental cues and behaviorally relevant goal locations (i.e., the target objects). Such a representation can then be indexed from any starting location, allowing individuals to determine their current location and plan a novel, efficient path.

We tested this capacity within runs by requiring participants to navigate to each object location three times (**Figure 1B**), each from a different starting position (**Figure S2, Panel A**). Starting positions were never proximal to the target, and as depicted in **Figure 1E-G** (still frame images of the participant view at the bottom), participants’ limited field of view was typically restricted to nearby environmental cues rather than the global layout. As a result, successful navigation required using local environmental information, which was not directly adjacent to the goal, to infer one’s position within the broader space and to plan a route based on the relationship between the current location and the target.

Under a map-based strategy, participants can use integrated spatial relationships of the global environment to generate efficient trajectories, resulting in both smaller distance error and lower excess path. This strategy can be better understood by looking at **Figure 1B**. Here, participants are cued to find the teal object, which is located between the traffic cone and the navy mountain (true location depicted by “x”), but begin facing the yellow mountain on the opposite side of the arena (depicted by grey dot). If a participant has represented the spatial relationship among these cues, including the yellow and navy mountains, as well as the traffic cone and target object, they can determine their current location within the map, turn around, and navigate directly toward the goal, resulting in both lower excess path and less distance error. Such a pattern would reflect both specific object-location memories, as well as a globally integrated representation that enables efficient trajectories to those objects from different starting positions (i.e., across different trials within a run).

In contrast, under a local, item-based strategy, participants form specific object-location memories without constructing a globally integrated cognitive map. Although they can still reach the correct location, they tend to do so through indirect, search-like trajectories. Following the previous example, rather than indexing the map to infer the correct direction to move toward, participants may only have knowledge that the teal object is near the navy mountain and/or the traffic cone. Thus, upon beginning at the yellow mountain on the opposite side of the arena, the item-based strategy may not support inference of efficient paths from the current location. Instead, participants must actively search the environment for the traffic cone or navy mountain via trial-and-error movements, resulting in less direct and more variable paths. Importantly, despite such inefficiency, the final distance error between the participant’s remembered location and the object’s true location can remain minimal. This manipulation therefore dissociates two forms of spatial knowledge: memory precision for individual object-location associations (scaled accuracy) and the ability to form integrated cognitive maps and flexibly express them via direct paths (excess path).

##### Updating spatial representations across environmental transformations

A second hallmark of cognitive maps is the ability to update them when they no longer support accurate behavior. That is, when the relationship between the internal map and the external environment is disrupted, individuals must determine whether the existing map is still applicable or whether a new representation is required. The ability to update the map, versus forming a new one, requires detecting structural invariance in spatial relationships despite surface-level changes. This updating process allows the map to be transformed and re-expressed, supporting rapid adaptation in response to the changing conditions.

To test this capacity, we examined whether participants could recognize that the relative positions of objects and the proximal landmark remained constant, even when we systematically altered the distal reference frame. Specifically, across runs, we rotated the distal cues by 90° or 180° while keeping object locations and the proximal landmark fixed (**Figure 1D-G**). Thus, this manipulation preserved the underlying spatial configuration (i.e., coordinate locations of the objects), but altered its alignment with the distal reference frame, allowing us to test whether participants could detect this invariance and realign their internal representation accordingly.

A map-based strategy should allow participants to recognize that object positions are unchanged within a common coordinate system and to mentally rotate their internal representation to align with the new distal frame. For example, as shown in **Figure 1F** (“Map-based interpretation”), participants can preserve their internal coordinate map (i.e., white dashed circle among objects) by rotating the distal mountain frame 90° to the left between Run 1 and Run 2 (i.e., pink mountain is now north), such that the objects retain their relative spatial positions even though their nearest distal mountain cues change (e.g., the pink object shifts from the yellow to the teal mountain). Behaviorally, this strategy should produce brief object-location errors immediately following rotation (i.e., first trial of Run 2), followed by rapid recovery and improvements in scaled accuracy and excess path values across subsequent trials and runs (i.e., first trial of Run 3), consistent with successful updating of the cognitive map. To further probe sensitivity to changing spatial structure, we alternated the lateral positions of the two distal objects across runs (e.g., the pink monster shifting from the left to the right mountain side; **Figure 1E-G**), maintaining global geometry while varying local contingencies.

In contrast, under a local, item-based learning strategy, participants encode objects individually by associating them only with their nearest cues. Each rotation therefore renders the environment effectively new. For example, if participants relied on item-based learning in Run 1, the orange and pink monsters in **Figure 1E**, originally adjacent to specific mountains, would appear to occupy entirely different positions in Run 2 (**Figure 1F**; “Item-based interpretation”). In coordinate terms **(Figure 1F**; “Item-based interpretation”), this strategy would require learning new object-location coordinates for each configuration change, each run. Behaviorally, such a strategy would manifest as memory improvement across repetitions within runs but little or no improvement across runs, reflecting repeated re-learning rather than updating. An intermediate profile, expected in adolescence, would show gradual improvements across runs, indicating partial updating of spatial relationships, but without the rapid, one-trial gains characteristic of fully integrated map-based updating (i.e., the first trial of Run 3 and Run 4, see below).

##### Generalization to new objects with the same spatial structure

A third property of cognitive maps is the ability to generalize existing spatial knowledge to new content (Behrens et al., 2018). Whereas the rotation manipulation in Runs 1-3 tested whether participants could update an existing object configuration to new spatial orientations, the second half of the task (Runs 4-6) examined a more abstract form of generalization: applying a previously learned spatial map to a new set of objects (**Figure 1G**). If participants have formed a holistic, map-like representation of spatial structure, they should generalize knowledge to the new objects, resulting in improved performance relative to initial learning (i.e., Run 1, Trial 1 vs. Run 4, Trial 1). In contrast, a purely item-based strategy would require re-learning object-location associations, yielding no such benefit.

##### Summary of task manipulations and behavioral predictions

Together, these manipulations (**Figure 1D**) allowed us to dissociate behavioral signatures of local, item-based spatial learning from the formation and flexible expression of a global, integrated cognitive map. The task required participants to learn multiple object locations, navigate to those locations from novel starting positions, update their representations when the distal reference frame changed, and generalize the learned spatial structure to a new set of objects. These manipulations therefore probed three complementary properties of cognitive maps across the task: flexible navigation, updating, and generalization.

The behavioral predictions followed from the degree to which spatial information was integrated across experiences. Limited integration across time and space should lead to a more local, item-based strategy, in which participants learn individual object-location associations but not their relationships. Under this profile, participants are expected to achieve relatively precise object-location memories, resulting in above-chance scaled accuracy, but to navigate to those object locations inefficiently, as evidenced by greater excess path. Such a strategy should also result in little transfer across changing task conditions (i.e., across runs), with improvements occurring primarily across repeated exposures within runs, at least for scaled accuracy.

In contrast, a mature, globally integrated representation should capture both individual object locations and the spatial relationships among those locations and environmental cues. Such a representation should support both precise object-location memory and efficient navigation from novel starting positions. Moreover, an integrated representation should enable participants to flexibly re-express the spatial representation when the distal reference frame changes, and to generalize the learned spatial structure to new object sets. We therefore expected map-based navigation to result in a pattern of behavioral change across runs, beginning with the initial detection of a mismatch between expected and experienced spatial relationships (Run 2), followed by updating of the representation, and leading to eventual stabilization.

First, when the environment changed unexpectedly at the beginning of Run 2, we expected an initial performance cost. Specifically, on the first trial of Run 2, participants should initially navigate toward the object locations learned in Run 1, reflecting the continued expression of the existing spatial representation despite the changed distal reference frame. The discrepancy between the predicted object locations and the newly experienced object locations should therefore generate a prediction error, signaling that the existing representation should be updated (or a new one formed). If participants subsequently update and realign this representation to the new spatial reference frame, they should show savings when the environment is then transformed again in Run 3. Accordingly, we compared performance on the first trial of Run 3 with the first trial of Run 1. Better performance on Run 3 relative to Run 1 would indicate that the relational structure acquired during the previous runs was re-expressed to facilitate new learning.

Whereas this trial-level comparison tested for rapid updating following initial environmental change, across-run analyses further tested whether the resulting spatial representation stabilized over subsequent experience. We therefore expected across-run improvements to occur primarily from Runs 1 through 3, as participants detected the prediction error induced by the unexpected environmental change in Run 2 and subsequently updated and realigned their spatial representation. Once this structure had been acquired, we did not expect further improvements across subsequent runs. Instead, performance was expected to remain relatively stable from Run 3 through Run 6, including following the introduction of new objects in Run 4, reflecting the continued re-expression of the learned spatial structure. Accordingly, we specifically expected differences between Run 1 and Run 4, but no differences between Run 3 with each of the generalization runs (Run 4, Run 5, or Run 6).

At the same time, and more broadly, we expected developmental differences in performance following each environmental change, such that older participants should show greater improvements relative to younger participants. Although we predicted that performance would stabilize following Run 2 for adults, we reasoned that the lower likelihood of extracting and representing the global relationship structure in younger participants may lead to disproportionate difficulty later in the task, as both the objects and distal reference frame change. As such, we hypothesized that older participants would show greater improvements relative to younger participants across Runs 2 through 6, and that age x run interactions might be possible. Thus, taken together, mature map-based navigation should be reflected in improvements in both scaled accuracy and decreased excess path across Runs 1 through 3, followed by relatively stable performance across subsequent runs, and pronounced performance gains relative to younger participants following continued environmental changes.

##### Temporal autocorrelation as an index of integrated representations

Importantly, these behavioral task manipulations also provided a framework for testing our central neural hypothesis: that developmental increases in anterior hippocampal temporal autocorrelation will enable increasingly integrated representations that link individual object locations into a holistic map of the environment, spanning individual trials, phases (learning versus test), and vantage points. As described above, participants navigated to multiple objects repeatedly within each run, from different starting positions, providing opportunities to integrate individual object-location memories into a representation of the broader environment. Moreover, although the specific object-location associations changed across runs, the underlying spatial configuration was preserved. Thus, successful navigation required participants to maintain this common spatial representation and re-express it to guide behavior across distinct experiences.

Temporal autocorrelation—which indexes the degree of representational stability over time—therefore provided a measure of this hypothesized integrative computation. If individual navigational experiences, distributed across time, are linked within a shared cognitive map, then neural representations should remain similar across otherwise distinct events—including across trials involving different objects and across different phases (i.e., learning versus test). Greater run-level temporal autocorrelation would therefore be consistent with the maintenance and continued expression of a common representation, rather than the formation of individual representations for each object or experience. We therefore quantified run-level autocorrelation as an index of the stability and continued expression of an integrated spatial representation and predicted that its relationship with navigation behavior would increase with development.

Importantly, by estimating temporal autocorrelation separately for each run, as discussed in the statistical approach section later, we could further model the association between run-level temporal autocorrelation and overall behavior on the corresponding run, within participants. This approach allowed us to evaluate whether the association between individual run-level hippocampal autocorrelation and behavioral performance was evident across each of the six task runs, despite changes in task contingencies. The approach was thus consistent with our broader theoretical question of whether stable hippocampal representations support behavior across changing task contingences that unfold over the course of the experiment.

In addition to examining the association between temporal autocorrelation and overall run performance, which we hypothesized for adults, we also examined whether run-level temporal autocorrelation was related to improvements across repetitions within individual runs. As noted above, our primary run-level autocorrelation/performance analyses stem from the framework that participants can form and re-express the same structured representation when the environment changes across runs. In other words, by flexibly re-expressing the same structured representation, participants may rapidly learn the new associations on the first trial of each new run, leading to more stable behavioral performance across repetitions within a run. However, if formation and re-expression of this structured representation is not developmentally possible— particularly in children, who may rely more on run-specific, locally formed representations—then overall performance on individual runs may be less sensitive to detecting brain-behavior associations. Instead, within each run, temporal autocorrelation may track the strengthening of a representation as participants strengthen individual associative memories over repeated test trials. Because each target object and location was tested three times within a run, the task design allowed us to quantify a trial-by-trial improvement score for scaled accuracy. Under this alternative framework, greater run-level temporal autocorrelation should be associated with repetition-related gains in accuracy, reflecting the stabilization or strengthening of the representation formed within that run, even if that association is not observed for overall performance on the run.

##### Dissociable predictions for how scaled accuracy and excess path relate to temporal autocorrelation

We next considered how these distinct forms of hippocampal representation—local versus global—should relate to the two behavioral measures, and how these relationships should differ across development. The critical distinction was that scaled accuracy and excess path place different demands on integrated representations. Scaled accuracy indexes the precision of individual object-location memories and therefore can be supported by either a local, item-based representation or a more globally integrated map. Excess path, in contrast, provides an index of map-based navigation, because efficient trajectories from novel starting positions require participants to use relationships among objects and environmental cues to determine their position within the broader environment and plan a direct route. These theoretical distinctions thus yielded dissociable predictions for the association between hippocampal autocorrelation with scaled accuracy versus excess path.

For scaled accuracy, which indexes the precision of object-location memory, we predicted complementary contributions along the hippocampal long axis. Posterior hippocampal autocorrelation may support stable, precise representations of object locations (Doeller & Burgess, 2008; Nadel et al., 2013), whereas anterior hippocampal autocorrelation should support the integration of those individual representations into a common spatial representation that can be maintained, updated, and re-expressed across experiences. Accordingly, we expected greater run-level temporal autocorrelation to relate to better scaled accuracy performance across both hippocampal subregions. We further expected this joint anterior-posterior profile to emerge most clearly in adults, whose behavior would show the strongest evidence of integrated representation that supports both precise object-location memory and map-based updating and generalization (**Figure 1F**; left arena).

We further predicted that the relationship between hippocampal temporal autocorrelation and scaled accuracy would strengthen with age, which could arise in one of two ways. Our primary hypothesis was that this age-related increase would be greater in anterior than posterior hippocampus. Such a pattern would suggest that anterior hippocampal representations become disproportionately more predictive of accurate object-location memory across development, consistent with an increasing role for integrating individual locations into a broader spatial structure. This hypothesis is consistent with prior work showing that anterior hippocampal development is particularly important for behavioral performance on tasks requiring the representation of associative structure across temporally separated experiences (Schlichting et al., 2016).

Alternatively, similar age-related increases in the relationship between temporal autocorrelation and scaled accuracy across anterior and posterior hippocampus would suggest parallel developmental increases in the behavioral relevance of both integrated and individual object-location representations. Under this account, successful learning of the task’s associative structure may support not only the formation of an integrated cognitive map, but also the stabilization and increasing precision of the individual object-location associations within that broader structure. In other words, if stable posterior hippocampal representations depend in part on successfully learning and integrating individual locations into a common spatial representation, anterior and posterior hippocampal representations may therefore become increasingly predictive of scaled accuracy in parallel across development. This account is also consistent with prior work showing age-related increases in posterior hippocampal granularity that predict detailed memory, suggesting that the behavioral relevance of posterior hippocampal coding may likewise strengthen with development (Callaghan et al., 2021).

A third and non-mutually exclusive possibility is that children may show little relationship between run-level autocorrelation and scaled accuracy because their improvements in object-location memory are driven primarily by incremental learning across repetitions rather than by stable expression of a run-level, global representation. To test this possibility, we examined whether repetition-related increases in hippocampal temporal autocorrelation track repetition-related increases in scaled accuracy within runs. A positive relationship would indicate that behavioral improvements within runs were accompanied by strengthening of individual object-location representations across repeated experiences, consistent with an item-specific learning strategy, which we predicted to be predominant in posterior hippocampus.

Finally, in contrast to scaled accuracy, excess path provided a more stringent test of the contribution of anterior hippocampal representations to map-based navigation. Because efficient trajectories require participants to locate themselves within the global environment and plan routes through that space, we predicted that excess path would be selectively associated with anterior hippocampal run-level autocorrelation, but not posterior hippocampal autocorrelation. We expected this relationship to be strongest in adults, for whom integrated spatial representations should be consistently deployed to guide navigation. In children (and possibly adolescents), weaker or absent associations would instead suggest that efficient navigation from novel starting positions is not yet consistently supported by a stable, globally integrated representation. Thus, whereas scaled accuracy should reflect complementary contributions of posterior and anterior hippocampus, excess path provides a more selective behavioral signature of the anterior hippocampal contribution to flexible, map-based navigation.

### MR image acquisition

Whole-brain imaging data was aggregated from three institutions with four separate scanning protocols (**Table S1**). HBN data was collected at the Rutgers University Brain Imaging Center with a Siemens Trio Tim 3T scanner and at CitiGroup Cornell Brain Imaging Center with a Siemens Prisma 3T scanner. NKI-RS imaging data was collected using a Siemens Trio Tim 3T scanner and UT Austin imaging data was collected using a Siemens Skyra 3T scanner. Across sites, imaging data included high-resolution T1-weighted images for co-registration and parcellation and functional T2*-weighted images either during rest or task. Fieldmaps were acquired prior to functional runs for the HBN and UT Austin samples to correct for magnetic field distortions. As described below (***Experimental design and statistical analy*ses**), we accounted for differences in acquisition parameters in all statistical models that included data from multiple sites.

### MR image preprocessing

Resting-state and task-based images were both preprocessed and analyzed based on the same preprocessing pipeline (Morton et al., 2021) and established approach for modeling motion artifacts in resting-state (Ciric et al., 2018) using FSL 6.0.4 (FMRIB’s Software Library, http://www.fmrib.ox.ac.uk/fsl), AFNI 16.2.0 (AFNI; Cox, 1996) and Advanced Normalization Tools 2.1.0 (ANTs) (Avants et al., 2011). Structural images were corrected for bias field with N4BiasFieldCorection and automatically segmented into cortical and subcortical areas using Freesurfer 6.0.1. All participants provided high-quality T1-weighted anatomical images, indicated by successful segmentation. Anatomical images were then registered to FSL’s 1mm MNI template image using ANTs.

Functional images were motion-corrected through alignment to the center volume using MCFLIRT with spine interpolation, computing framewise displacement (FD) and DVARS for each volume. To rigorously control for motion, which is known to cause artifacts in resting-state and developmental samples, runs were excluded from further analyses if more than 25% of their total volumes exceeded a FD threshold of 0.3 mm. Because the resting-state data was used to examine more stable, behavior-independent differences in intrinsic functional organization, and because participants provided only one (HBN, UT Austin) or two (NKI-RS) scanning runs depending on site, all resting-state runs meeting motion thresholds were included. For the task-based sample, which was designed to link neural maturation to behavior, participants who did not complete at least four out of six runs within predefined motion thresholds were excluded to ensure adequate sampling for the reliable estimation of brain-behavior effects (11 children, 1 adult excluded). For both studies, volumes exceeding motion thresholds were identified and later censored during preprocessing.

Next, functional scans were unwarped via boundary-based registration using an adapted version of FSL’s epi_reg (Greve & Fischl, 2009). Intensity-based registration was then implemented with ANTs to correct misalignments between functional and anatomical images. Non-brain tissue was removed based on automated brain masks extracted from Freesurfer in each participant’s functional space. These unwarped functional scans were then registered to the T1 anatomical images and transformed into anatomical space using trilinear interpolation via ANTs and FSL’s *flirt*. Bias field correction was implemented by dividing the functional timeseries by a single bias field image estimated from the average. Consistent with previous resting-state work (Ciric et al., 2018), the functional data was not smoothed.

Finally, all functional images were processed for motion using the motion parameters derived above. These motion parameters were applied using an established resting-state approach to yield whole-brain timeseries data for each participant (Ciric et al., 2018). We implemented a 32-parameter regression model including continuous timeseries regressors for cerebral spinal fluid, white matter, and six motion parameters and their respective temporal derivatives and squares extracted using FSL’s MCFLIRT and *fslmeants.* Timeseries data was extracted for each participant in native space using AFNI’s *3dTproject* to create the model with the pre-processed BOLD data, motion parameters, and a high-pass temporal filter (s = 64s; Gaussian-weighted least-squares straight line fitting). For all high-motion volumes flagged during spatial pre-processing, linear interpolation was used to replace each voxel’s value based on neighboring non-censored TRs. Each timeseries was then masked with anterior and posterior hippocampal masks using *fslmeants*, ultimately producing a voxel-by-TR matrix for each run within each participant in each region of interest in native functional space.

### Regions of interest (ROIs)

Primary analyses were conducted in *a priori* bilateral anatomical hippocampus, from which separate anterior and posterior subregion ROIs were defined. Following past developmental protocols (e.g., Riggins et al., 2015), the hippocampus was divided based on uncal location to delineate anterior hippocampus (before uncus) and posterior hippocampus (after uncus; including body and posterior portions of hippocampus). Following past work (Brunec et al., 2018; Callaghan et al., 2021), an additional anatomical ROI corresponding to bilateral parahippocampal cortex was defined as a control region to interrogate whether changes in temporal autocorrelation were specific to hippocampus, or rather, to medial temporal lobe (MTL) cortex more broadly. All ROI masks were drawn manually on the FSL 1mm MNI template. They were then down-sampled to the 2mm MNI template with FSL’s *flirt* function and transformed into each participant’s native space for further analysis using ANTs.

### Analytical approach

#### Experimental design and statistical analyses

Statistical analyses were conducted with custom R code (version 2026.05.0), which is available at https://osf.io/g7qne. Continuous outcome variables were z-scored prior to model fitting, and categorical predictors were specified as factors (using R’s default treatment coding). Linear mixed-effects models were fit using the lme4 package, assuming Gaussian residual errors after accounting for fixed and random effects (Bates et al., 2015). Random intercepts were included to account for repeated observations within participants and other sources of non-independence introduced by the repeated runs of scanning in our study design. Critically, acquisition site was included as a covariate in all resting-state analyses involving data collected across multiple sites to account for variance in acquisition parameters (**Table S1**) when assessing developmental differences.

Neural models were specified with fixed effects for age group, hippocampal subregion, the hypothesis-specific performance predictor (i.e., scaled accuracy or excess path), and corresponding interactions, while controlling for hemisphere, motion, image quality, anatomical, acquisition-related, and experiment-specific covariates. Neural models included nested random intercepts for participant and run (1|subject/run) to control for variability across scans and participants, but were not analyzed by run and therefore did not include fixed effects of run. Rotation order across individual runs was additionally included as a run-level covariate in all task-based models to account for potential effects of the specific configuration presented, termed “run_configuration” in models.

Behavioral models were specified analogously, including random intercepts for participant (1|subject), but excluded hippocampal subregion effects. Run-level effects were excluded from behavioral models examining aggregate (across-run) behavior, while fixed effects of run were included in models that directly interrogated pairwise differences between individual runs.

Across both experiments, fixed effects and interaction terms were specified based on our *a priori* hypotheses about developmental differences in anterior hippocampal temporal autocorrelation, and its relationship to behavior. To that end, in both experiments, fixed effects and interactions were first specified between age and subregion, testing whether the relationship between temporal autocorrelation and age differed in a subregion-specific manner (i.e., a significant age x subregion interaction; we predicted age-related increase in anterior hippocampus). For the task-based experiment, fixed effects and interactions were then further specified among performance, age group, and subregion based on *a priori* hypotheses that either: (1) the relationship between temporal autocorrelation and performance would differ across age in a subregion-specific manner (i.e., a significant age × behavior × subregion interaction; we predicted age-related increase in the association between performance and anterior hippocampus, for both scaled accuracy and excess path), or (2) the relationship between temporal autocorrelation and performance would differ across age groups, but these developmental differences would appear in both subregions (i.e., a significant age × behavior interaction; we predicted age-related increase in the association between performance and temporal autocorrelation across the whole hippocampus; an alternative prediction for scaled accuracy, see above). Accordingly, performance models first evaluated the hypothesized higher-order (three-way) interaction. In cases when the three-way interaction was not significant, it was removed and only lower-order interactions that were specified *a priori* were evaluated.

For both neural and behavioral models, statistical significance was evaluated at an alpha level of *p* < .05, and all reported statistics were two-tailed. In addition, planned pairwise comparisons were specified *a priori* to test primary experimental hypotheses about differences in behavior across runs and are described in detail alongside their corresponding analysis in the Results, while all pairwise contrasts estimated by the models are reported in the OSF repository (https://osf.io/g7qne). Pairwise comparisons were extracted via an estimated marginal means approach with Bonferroni correction using the *emmeans* package in R.

To facilitate transparency and clarity in reporting the more complex neural models, which included a greater number of factors, covariates, and interactions terms, analysis-specific model equations and complete model parameters for the neural models are reported in the supplementary materials. Due to the extensive number of behavioral models, and their comparatively more straightforward nature, behavior-only models are described in the main text without corresponding tables; complete model specifications and statistics are available on the OSF repository (https://osf.io/g7qne).

#### Quantifying temporal autocorrelation

Temporal autocorrelation was calculated separately for each ROI (posterior hippocampus, anterior hippocampus, parahippocampal cortex) within each functional run. For each timepoint (TR), a vector containing the activity of all voxels within an ROI (**Figure 2A-2B**) was extracted using *fslmeants*. Temporal autocorrelation was then computed as the Pearson correlation between vectors from adjacent TRs (e.g., TR_2_ with TR_1_), reflecting a vector of all the correlation values of all of the voxels to themselves. These comparisons yielded one similarity measure per ROI per pair of adjacent timepoints (i.e., one fewer than the total number of TRs).

**Figure 2.**
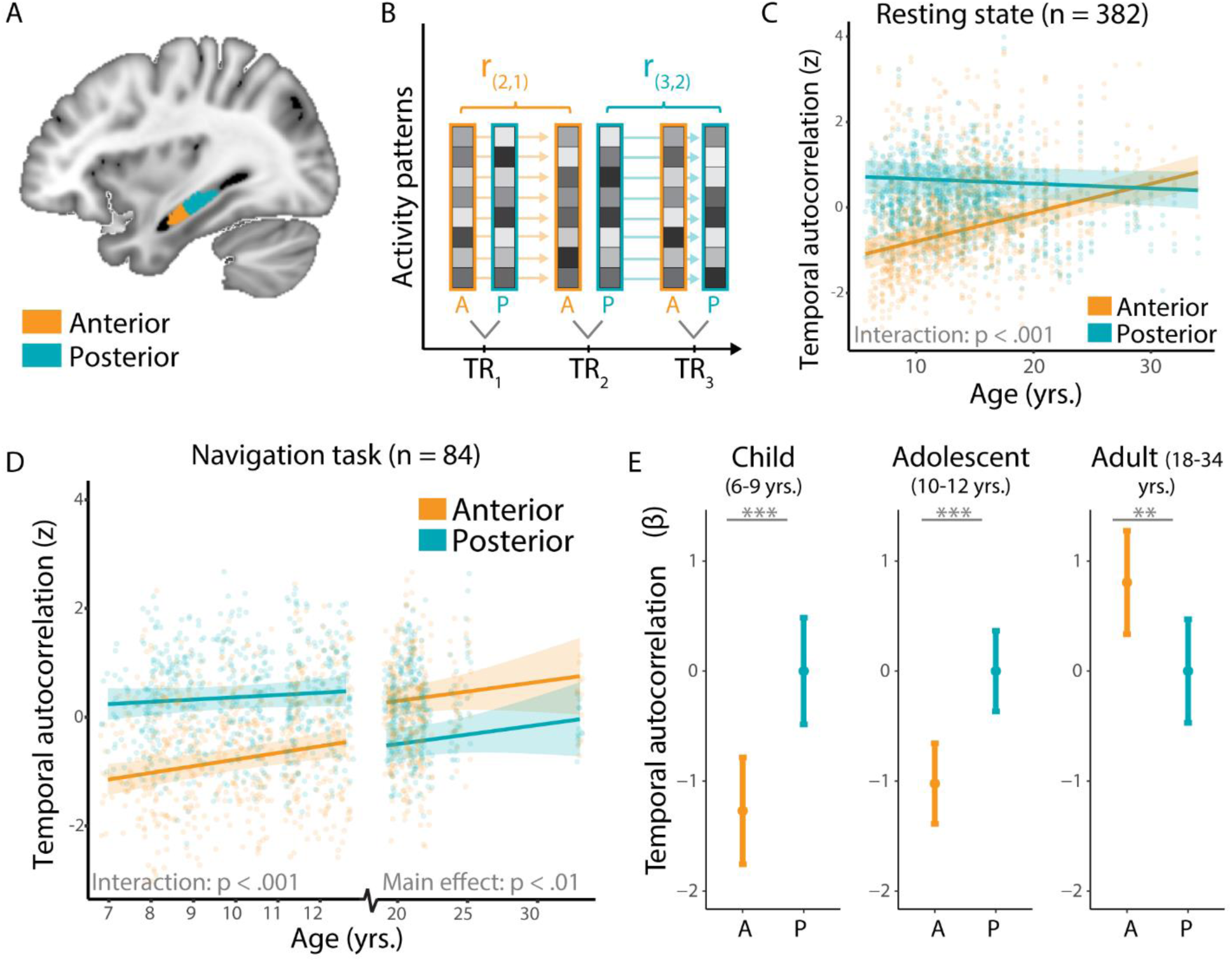
Analytical approach to quantifying temporal autocorrelation by hippocampal subregion. **A**) Hippocampal masks were drawn manually on the 1mm MNI template based on uncal location (anterior = before uncus, posterior = after uncus) and transformed into each participant’s native space using ANTS for all analyses. **B**) Temporal autocorrelation is calculated by deriving Pearson correlations between all neural activity at contiguous TRs within each hippocampal subregion. **C**) Age x subregion interaction demonstrated in a large, continuous resting state sample from age 5-34 years. Trend lines and 95% confidence intervals reflect mixed-effects model estimates for age differences in autocorrelation within each subregion. Points indicate individual (repeated) observations, with four per run of resting-state data, corresponding to each unique subregion (anterior, posterior) and hemisphere (left, right) combination. Temporal autocorrelation is plotted as a z-score, with a z-score of 0 (i.e., mean autocorrelation) corresponding to Fisher’s Z of 1.67 (SD = 0.31). **D**) Spatial navigation task highlights age and subregion differences in temporal autocorrelation. As in the resting-state sample, we identify an age x subregion interaction in the developmental group (6-12 years) indicating that anterior hippocampal autocorrelation increases at a greater rate during development than posterior hippocampal autocorrelation. Further, adults (18-33 years) show greater temporal autocorrelation in anterior relative to posterior hippocampus, invariant of age. Trend lines and 95% confidence intervals depict mixed-effects model estimates for age differences in autocorrelation (z-scored) within each subregion and are separately modeled in the developmental and adult groups. Points indicate individual (repeated) observations, resulting in four points per run corresponding to each subregion (anterior, posterior) and hemisphere (left, right). Temporal autocorrelation is plotted as a z-score, with a z-score of 0 (i.e., mean autocorrelation) corresponding to Fisher’s Z of 2.03 (SD = 0.27). **E**) Coefficient estimates extracted from task-based mixed-effects models illustrate subregion differences in autocorrelation by age group, demonstrating group estimate of anterior relative to posterior autocorrelation. Values and significance tests are extracted directly from mixed-effects models. Points reflect model coefficient estimates for anterior autocorrelation relative to posterior (reference level = 0) and error bars depict standard error. Coefficient estimates indicate that adults demonstrated significantly greater temporal autocorrelation in anterior hippocampus than children and adolescents, and adolescents demonstrate significantly greater anterior autocorrelation than children. *p < .05. **p < .01, ***p < .001. Participants provided four individual observations per run (anterior and posterior hippocampus in left and right hemisphere), resulting in 2504 measurements represented in **Panel C** (resting-state sample) and 1888 data points represented in **Panel D** (task-based sample).

Correlation coefficients were normalized using Fisher z transformations before statistical analyses. Because our primary hypotheses concerned developmental differences between anterior and posterior hippocampus, autocorrelation values were summarized at the level of anatomically defined hippocampal subregions rather than treated as single-voxel outcome measures. Specifically, Fisher z-transformed autocorrelation values were averaged across voxels within each ROI, run, and hemisphere, yielding one estimate for each anterior and posterior hippocampal subregion in each hemisphere per run.

This ROI-level approach differs from single-voxel analyses that are optimized for mapping fine-grained temporal gradients or local heterogeneity within hippocampal subregions. Our goal was instead to test whether the overall temporal stability of activity patterns within anterior versus posterior hippocampus changes with age and relates to navigation behavior. Aggregating across voxels therefore provides a more reliable subregional estimate, reduces sensitivity to voxel-specific noise and anatomical variability in a large developmental sample, and aligns with prior developmental work comparing hippocampal subregions in volume and activation (e.g., Demaster et al., 2016; DeMaster & Ghetti, 2013; Sastre et al., 2016; Schlichting et al., 2016). This approach allowed us to test whether anterior hippocampal autocorrelation shows a more protracted developmental trajectory than posterior hippocampal autocorrelation while maintaining comparability with earlier studies of hippocampal organization (Ghetti & Bunge, 2012; Murty et al., 2016).

This same general procedure was applied across all samples (resting-state and task-based) with one exception. Repetition time (TR) varied across acquisition sites, with the HBN protocol using a shorter TR (0.8s) than the others (1.4-2.0s; **Table S1**). To ensure a comparable temporal lag between measurements, autocorrelation in the HBN resting-state data was computed across every other TR, corresponding to a 1.6 s lag. Autocorrelation values were z-scored within each site to further account for protocol differences. Finally, as noted above, acquisition site was included as a covariate in all resting-state analyses to further account for differences in acquisition parameters.

#### Modeling temporal autocorrelation as a function of age and subregion

A key question in both the resting-state and task-based samples was whether temporal autocorrelation varied as a function of hippocampal subregion (**Figure 2A**) and age, with greater age-related increases in anterior, but not posterior, hippocampal autocorrelation. Models were estimated using hierarchical mixed-effects regression with a nested run-within-participant random intercept. For our primary hippocampal analyses, subregion (anterior vs. posterior) was included as a fixed effect, resulting in four autocorrelation outcome measures per run per participant (anterior-left, anterior-right, posterior-left, posterior-right). This structure allowed temporal autocorrelation to be modeled separately for each subregion within each run while accounting for non-independence of observations nested within individuals. Image-quality covariates were included in all models (see **Covariates and model comparisons**), calculated separately for each participant, run, and ROI.

We further tested whether subregional differences in hippocampal temporal autocorrelation differed as a function of age. To that end, in the resting-state sample, which was sampled continuously from 5-34 years, age was treated as a continuous predictor in years and months. In the task-based sample, which was discontinuous (ages 6-12 years and 18-34 years), we first implemented the same continuous analysis as in the resting-state approach (separately for the developmental and adult samples), providing a replication and extension to an independent, task-based context. As noted above (see **Participants**), participants were also grouped into three *a priori* bins comprising children (6-9 years), early adolescents (10-12 years), and adults (18-34 years) and thus additionally modeled as a categorical predictor, enabling tests of hypothesized associations between subregional hippocampal autocorrelation at theoretically motivated age windows.

In addition to our primary models assessing linear age effects, we conducted exploratory analyses to examine potential non-linear age patterns. Specifically, we re-ran the primary analyses including quadratic and cubic polynomial age terms. No significant non-linear effects of age were observed in either the resting-state or task-based samples (ps > .05), and none of the polynomial terms interacted with subregion. Therefore, non-linear models were not considered further.

#### Covariates and model comparisons

All neural models included covariates to account for potential sources of variance related to image quality and acquisition. As noted above, for all resting-state analyses, acquisition site was included as a covariate to account for differences in acquisition parameters across the aggregated datasets. Temporal-signal-to-noise ratio (tSNR) was calculated as the mean signal intensity divided by its standard deviation and was included to control for signal quality over time. Two motion metrics were also included: mean absolute translation (overall motion across the scan) and mean FD (variability in motion between successive timepoints).

To further account for other acquisition and anatomical differences, models also included the number of voxels within each ROI mask, the number of TRs per scan, and the average distance between voxels in the x-, y-, and z-axes within each mask, following prior work (Brunec et al., 2018). These quality metrics were computed separately for each participant, run, ROI, and hemisphere. Control analyses also ensured that biological sex and general cognitive ability did not account for differences in autocorrelation within the navigation sample, indexed using the vocabulary subtest of the WASI-II (Wechsler, 2011).

Because several covariates captured similar measures of motion or image quality, we conducted a model comparison approach to assess potential collinearity. Specifically, we iteratively removed nuisance covariates and compared reduced models to the full model using likelihood ratio tests. Notably, removing motion variables did not change primary effects of interest, and no reduced model significantly improved model fit (Chi-sqs. < 2.9, ps > 0.09). In many cases, removing covariates increased AIC, indicating improved model fit when quality metrics were retained. Therefore, all reported models include the full set of quality-related covariates.

#### Behavioral analyses

To assess developmental differences in navigation performance, we implemented linear mixed-effects models predicting scaled accuracy and excess path (outcome variables) from age group, run and their interaction (fixed effects), with participant-level random intercepts to account for individual variability. All model coefficients reported in the text were extracted directly from the fitted models.

For each model, we first evaluated main effects of age group and run, as well as their interaction, by examining their corresponding coefficients (which are reported in the main text). Pairwise contrasts were subsequently examined to characterize significant omnibus effects. Although the fitted models generated estimates for all pairwise contrasts among levels of the categorical predictors, given the sheer number of comparisons conducted, our reporting focused on the most informative contrasts and those corresponding to our *a priori* hypotheses (e.g., Run 1 vs. Run 3). However, complete model statistics are available on the OSF repository. All pairwise comparisons were estimated using marginal means and Bonferroni correction.

We followed the same analytic approach for supplementary behavioral analyses. To further characterize patterns of behavioral performance across object conditions, we assessed scaled accuracy and excess path between proximal and distal trials using analogous mixed-effects models with the same structure and main and interaction effects for condition (proximal vs. distal) but with participant-level data derived from trials within each trial-type. Finally, we fit analogous models for navigation time, quantified separately for learning and test trials. These analyses served as control analyses for potential differences in motor abilities and to identify the number of fMRI measurements to include as covariates in the neural models, respectively.

#### Modeling temporal autocorrelation and performance associations

After interrogating developmental effects of subregional hippocampal autocorrelation, and behavioral performance, in our final set of analyses we further tested whether run-level temporal autocorrelation—our index of the stability and continued expression of an integrated representation across distinct experiences within a run—was associated with navigation performance. These analyses used an analogous hierarchical mixed-effects modeling framework to maintain the same random-effects structure and covariates as the primary neural models described above. Here, separate models were estimated for each primary behavioral measure: run-level scaled accuracy and run-level excess path. Estimating all effects within a single, fully adjusted model allowed behavioral associations to be estimated while accounting for all model terms and covariates, rather than through separate age-stratified analyses.

To test our *a priori* predictions, for both behavioral measures, we first fit a full three-way interaction model including age group, performance, and hippocampal subregion, as well as main effects of performance and all lower-order interactions. These models evaluated our primary hypotheses that developmental increases in the relationship between hippocampal temporal autocorrelation and navigation performance would be greater in anterior than posterior hippocampus. However, if the three-way interaction was not significant for scaled accuracy, it was removed and the reduced model was used to evaluate the alternative *a priori* hypothesis of two-way interactions, namely, increases in the association among performance and age irrespective of subregion.

Follow-up analyses were conducted only when interactions were significant. Specifically, when three-way interactions were observed, behavior-autocorrelation slopes were examined within each age group and subregion to evaluate how behavior-autocorrelation associations differed across development and subregion. Similarly, when both age group x performance and subregion x performance interactions were significant (even in the absence of a three-way interaction), the same follow-up approach was followed to further characterize the nature of these effects. Importantly, all follow-up analyses by age group were extracted from these full models rather than modeled within each age group separately. All reported associations, even those featuring a single follow-up age group, therefore reflect relations between hippocampal autocorrelation and behavior above and beyond age and other covariates.

Finally, in addition to the primary run-level analyses, we conducted complementary models testing whether run-level temporal autocorrelation was associated with within-run learning. To this end, we computed a within-run behavioral improvement score, wherein for each target object, we calculated the change in scaled accuracy between consecutive repetitions (Repetition 2 – Repetition 1 and Repetition 3 – Repetition 2) and then averaged these scores across the four target objects within each run. Positive values therefore reflected increasing navigation accuracy across repetitions within a run. We then associated these within-run improvement scores to run-level hippocampal autocorrelation using the same hierarchical mixed-effects framework as the primary analyses, retaining the same random-effects structure and neural and participant-level covariates. Importantly, these models also retained all effects identified in the primary brain-behavior analyses, including those with run-level scaled accuracy, thus ensuring that the within-run improvement term in the alternative model captured variation in repetition-related increases beyond variation in overall navigation performance.

## Results

### Age-related increases in anterior hippocampal autocorrelation at rest and task

We first examined developmental differences in hippocampal temporal autocorrelation in the resting-state sample. We observed an age × subregion interaction, such that temporal autocorrelation increased more steeply with age in anterior than in posterior hippocampus (*B* = 0.079, SE = 0.006, *t* = 12.774, *p* < .001; **Figure 2C**; **Table S2**). Subregion-specific follow-up analyses confirmed significant developmental increases in anterior hippocampus (*B* = 0.072, SE = 0.008, *t* = 8.857, *p* < .001) but not posterior hippocampus (*p* = 0.388; **Table S3**). These patterns were consistent across temporal lags up to 20 s (**Supplemental Materials: Hippocampal autocorrelation across varying temporal lags**) and were not observed in parahippocampal cortex (*p* = 0.930; **Table S4**), indicating a prolonged, region-specific maturation of anterior hippocampal temporal autocorrelation.

To characterize subregional differences in adulthood, we next restricted the analysis to adult participants. Anterior hippocampal autocorrelation was higher than posterior hippocampal autocorrelation among adults 18 to 35 years (*B* = 0.603, SE = 0.305, *t* = 1.979, *p* = 0.049), but was not significant when restricting the sample to adults aged 20-35 years, a relatively older subset whose null result is consistent with prior work examining a similarly aged adult sample (Brunec et al., 2018).

We observed similar developmental increases in anterior hippocampal temporal autocorrelation in the independent task-based fMRI sample, both when age was considered continuously (6-12 years; **Figure 2D**; **Table S5**; *B* = 0.079, SE = 0.018, *t* = 4.322, *p* < .001) and when binned into our *a priori* age groups (**Figure 2E**; **Table S6;** *p*s < .001). These analyses replicate stronger age-related increases in anterior relative to posterior hippocampus, with developmental increases observed specifically in anterior hippocampus (*B* = 0.118, SE = 0.033, *t* = 3.592, *p* < .001; **Table S7**) and no corresponding effects in posterior hippocampus (*p* = 0.240; **Table S7**) or parahippocampal cortex (*p*s > 0.123; **Table S8**). These effects were not associated with sex (*p*s > 0.593) or general cognitive ability (*ps* > 0.166; **Tables S9-S10**).

Further, within individual age groups, the relative strength of anterior versus posterior autocorrelation differed (**Figure 2E**): adults exhibited greater anterior than posterior autocorrelation (*B* = 0.784, SE = 0.240, *t* = 3.272, *p* = 0.001), consistent with prior task-based findings in young adults (Brunec et al., 2018), whereas children (*B* = −1.266, SE = 0.180, *t* = −7.023, *p* < .001) and adolescents (*B* = −1.018, SE = 0.187, *t* = 5.454, *p* < .001) exhibited the opposite. Having established age-related increases in anterior hippocampal temporal autocorrelation, we next assess spatial navigation behavior, followed by relations between performance and temporal autocorrelation.

### Developmental differences in spatial navigation behavior reveal a shift from local, item-based learning to cognitive map formation and expression

We next examined age-related differences in spatial navigation performance to test whether children, young adolescents, and adults differed in their learning profiles. Specifically, we tested whether they shifted from an item-specific, object-location strategy within runs to an across-run, cognitive map-based strategy that supports updating and generalization.

#### Initial learning and navigation efficiency

We first confirmed that all age groups successfully learned the initial object-location associations in Run 1, before any contingencies changed. Scaled accuracy indicated that all groups performed above chance across trials in Run 1 (one-sample t-test vs. 0; *t*s > 11.890, *ps* < .001; **Figure 3A**). Scaled accuracy also improved across trial repetitions within a run (trial 1 to 3), significantly for all age groups (main effect: F(1, 875) = 45.979, p < .001; children: *t*(875) = 2.011, *p* = .045; adolescents: *t*(875) = 2.405, *p* = .016; adults: *t*(875) = 5.581, *p* < .001), indicating that participants used trial-by-trial feedback to improve memory within a run. Age group comparisons on Run 1 further revealed developmental differences (main effect; F(2, 81) =7.609, p < .001). Adults were more accurate than children (*t*(81) = 3.863, p < .001), whereas adults and adolescents (*t*(81) = 2.400, *p* = 0.056) and adolescents and children (*t*(81) = 1.572; *p* = 0.359) did not differ significantly from one another.

**Figure 3.**
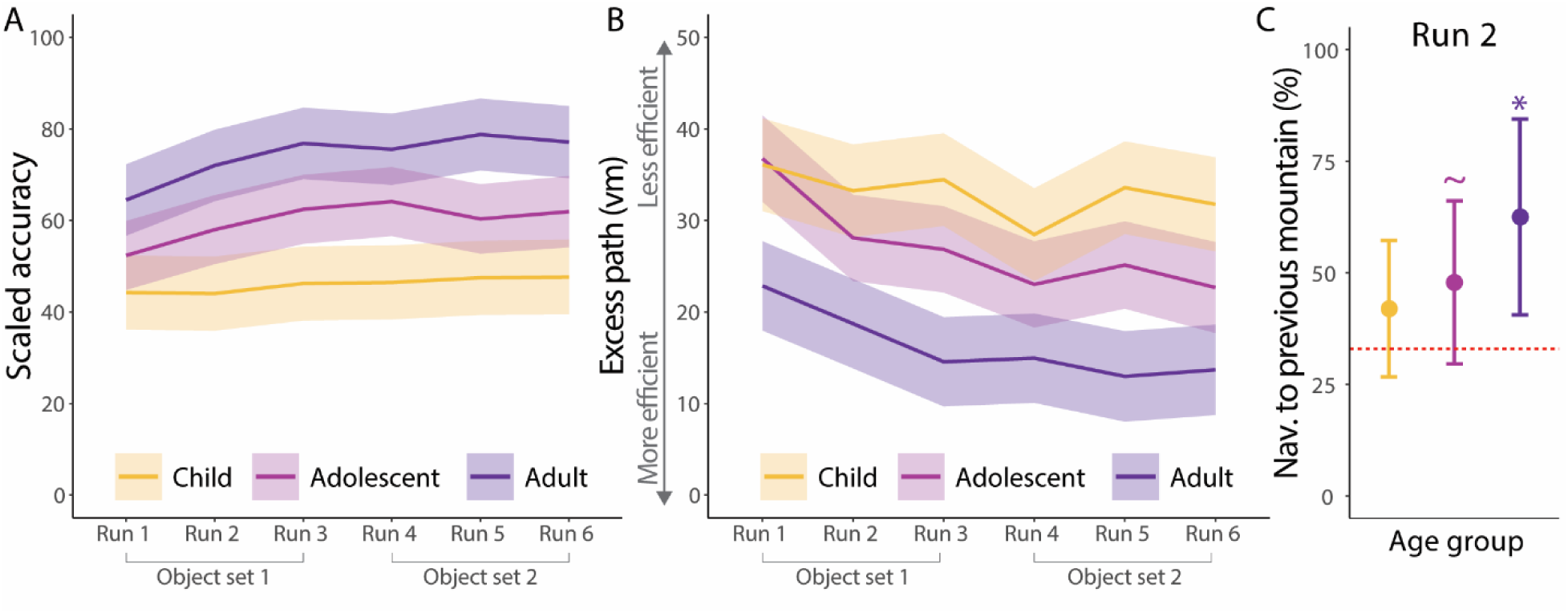
**A**) Run-level scaled accuracy by age group. Scaled accuracy increased with age, and adults and adolescents improved run-to-run while children did not. Lines reflect average scaled accuracy while shading reflects standard error of the mean (SEM). **B**) Excess path by run and age group. Excess path values decrease (i.e., efficiency improves) with age, and adults and adolescents become more efficient run-by-run from Runs 1 to 3. Lines reflect average excess path while shading reflects SEM. **C**) Percent of trials in which, when participants navigated to the incorrect mountain on the first repetition of Run 2, they navigated to the mountain from the previous run, reflecting a prediction error. Points and error bars reflect mean and confidence intervals derived via one-sided t-test (chance = 33%).

For excess path, there is no measure of chance performance. However, we again observed a main effect of trial (F(2, 302, = 7.874, p < .001), such that adults (*t*(864) = 5.360, *p* < .001) and adolescents (*t*(864) = 2.692, *p* = 0.007) showed reductions in excess path values from trial 1 to trial 3 within a run. Although children showed a similar pattern, the decrease across trials within a run did not reach significance (*t*(864) = 1.717, *p* = 0.090). As was observed for scaled accuracy, there were also differences across age groups (main effect: F(2, 80) = 8.934, p < .001), such that adults had lower excess path values than both adolescents (*t*(80) = −3.366, *p* = 0.004) and children (*t*(80) = −3.874, *p* < .001), while children and adolescents did not differ (*p* = 1.0).

Together, these results confirm that all age groups learned the initial object locations. Scaled accuracy improved across repetitions in all groups, and excess path values decreased across repetitions particularly for adolescents and adults. Adults also demonstrated lower excess path values, suggesting that although participants of all ages were able to form and retrieve precise location memories, adults flexibly navigated more efficient paths through the global environment, consistent with formation of initial integrated representations that may not have been accessible to other age groups.

#### Changing task contingencies reveal increasing developmental differences in navigation performance

Having established that all groups successfully learned the initial object-location associations in Run 1, we next examined whether performance remained reliably above chance as task contingencies changed. Confirming this pattern was critical before assessing developmental differences in how such performance was achieved. All groups continued to show above-chance accuracy as task contingencies changed, first through location rotations (two-sided t-test vs. 0: Runs 2-3; *t*s > 15.379, *ps <* .001) and then when new objects were introduced (two-sided t-test vs. 0: Runs 4-6; *t*s > 19.890, *ps* < .001).

Although all groups performed above chance, clear age-related differences emerged. In contrast to Run 1, during which only adults and children differed in scaled accuracy, across Runs 2 through 6 all age groups differed (main effect: F(2, 81) = 17.899, p < .001): adults were more accurate than both children and adolescents (adults > children: *t*(81) = 5.999, *p* < .001; adults > adolescents: *t*(81) = 3.058, *p* = 0.009), and adolescents were more accurate than children (*t*(81) = 3.094, *p* = 0.008; **Figure 3A**).

A similar pattern emerged for excess path (main effect: F(2, 28.750, p < .001): adults exhibited lower excess path than both younger groups (adults > adolescents: *t*(80) = −4.066, *p* < .001; adults > children: *t*(80) = −7.395, *p* < .001), replicating Run 1. In contrast to Run 1, here adolescents showed lower excess path than children (*t*(80) = −3.454, *p* = 0.003; **Figure 3B**). Thus, while all groups successfully learned the new associations, performance differed by age, with both location memory and efficiency improving with development.

#### Across-run improvements suggest greater use of prior spatial structure with age

The preceding analyses show that all age groups maintained above-chance scaled accuracy as task contingencies changed, but above-chance performance alone does not reveal whether participants used prior spatial structure or relearned locations within each run. We therefore next examined whether performance improved across runs. Such improvements would suggest updating or generalization of prior spatial knowledge.

To test our *a priori* prediction of across-run improvements from Run 1 through 3 in older participants, reflecting the updating and realignment of spatial representations following the initial change introduced in Run 2, we conducted planned Bonferroni-corrected pairwise comparisons. Scaled accuracy increased from Run 1 to Run 3 in both adults (*t*(397)=3.547, *p*=.006) and adolescents (*t*(397)=2.990, *p*=.044; **Figure 3A**). We further predicted that performance would stabilize thereafter, reflecting re-expression of the same integrated representation despite the introduction of new objects in Run 4 and continued rotation of the distal reference frame across all runs. Consistent with this prediction, performance did not differ between Run 3 and any of Runs 4-6 in either adults or adolescents (ts(398) < 1.118; *p*s = 1.0). A similar pattern was observed for excess path, with decreases in excess path from Run 1 to Run 3 in adults (*t*(392) = 3.162, *p* = 0.025) and adolescents (*t*(392) = 3.490, *p* = 0.008), and no further changes thereafter (Run 3 vs. Runs 4-6; ts(395) < 0.214; *p*s = 1.0).

In contrast, children showed no improvement across Runs 1-3 in either measure (*p*s = 1.0). Significant run x age group interactions further confirmed the pattern of developmental differences in across-run learning. Adults showed greater improvements than children in scaled accuracy from Runs 1 to 3 (*B* = 10.404, SE = 4.958, *t* = 2.099, *p* = .037), whereas adolescents showed a similar but non-significant difference (B = 8.106, SE = 4.878, t = 1.662, p = 0.098). For excess path, adolescents showed a greater decrease than children from Runs 1 to 3 (*B* = −7.364, SE = 3.578, *t* = −2.059, *p* = .041), with a comparable trend in adults (B = −6.663, SE = 3.608, t = −1.847, p = 0.067). These age differences were unlikely to reflect differences in basic motor abilities or exploratory tendencies, as navigation times to the same visible objects during initial learning trials did not differ across age groups (**Figure S3, Panel A**).

The developmental differences in initial learning also persisted following the change in object set on Run 4. Specifically, relative to children, adolescents improved more in scaled accuracy from Run 1 to Run 4 (*B* = 9.572, SE = 4.520, *t* = 2.122, *p* = .034) and adults showed a similar trend toward more improvement (B = 8.869, SE = 5.026, t = 1.765, p = 0.078). Notably, across-run improvements did not differ between adolescents and adults (*p*s > .197), suggesting broadly similar across-run learning gains in the older groups.

Converging evidence from condition-level analyses further supports this interpretation. Across runs, there was a main effect of condition on both scaled accuracy (**Figure S4: Panel A**) and excess path (**Figure S4: Panel B**), with all age groups showing better performance for proximal than distal objects. This pattern is consistent with the task manipulation, which altered object associations with the distal cues across runs while maintaining object associations with the proximal cue (**Figure 1E-G**). Critically, age interacted with this effect: older groups showed more comparable performance across the conditions than children. Specifically, whereas adolescents and adults showed similar scaled accuracy for proximal and distal trials, children performed significantly better on proximal than distal trials (**Figure S4, Panel A**). This pronounced proximal advantage suggests that children’s lower (**Figure 3A**), albeit above-chance, scaled accuracy may reflect greater difficulty learning the changing distal object-location associations, rather than a general propensity toward nonspecific spatial responding.

The pattern for excess path further distinguishes the developmental profiles of adolescents and adults. Although adolescents resembled adults in scaled accuracy, exhibiting similar performance across proximal and distal conditions (**Figure S4, Panel A**), their excess path values, like those of children, differed between conditions. Specifically, only adults showed comparable excess path values across conditions (**Figure S4, Panel B**). This dissociation suggests that adolescents show emerging sensitivity to changing contingencies in the distal condition, but do not yet leverage this information to support efficient, map-based navigation to the same extent as adults.

#### Rapid generalization reveals adult-specific map-based behavior

Although across-run improvements and condition effects suggest that adults and adolescents may have integrated information across runs, a more stringent test of generalization is performance on the first trial of a new run, when participants cannot rely on within-run feedback. We therefore examined behavior on the first trials of Runs 2 and 3 with the first trial of Run 1, providing a common baseline for assessing the initial cost of environmental change (Run 2; decreased accuracy) and subsequent savings following spatial updating (Run 3; increased accuracy and decreased excess path).

When object-location mappings were first rotated in Run 2, adults exhibited robust memory prediction errors, navigating to the original Run 1 mountain location at above-chance levels (*t*(15) = 2.360, *p* = .016; **Figure 3C**; distal objects only). Children showed no evidence of this effect (*t*(28) = 0.538, *p* = 0.298), and although adolescents exhibited a numerical trend in the same direction, it did not reach significance (*t*(22) = 1.392, *p* = 0.09). Adults also demonstrated rapid updating, showing higher scaled accuracy on the first trial of Run 3 compared to Run 1 (*t*(875)=5.625, *p* < .001; all objects), indicating that they used prior-run knowledge to rapidly update in the face of the rotated distal frame. Children again showed no evidence of this effect (*t*(875) = 1.305, p = 1.0), whereas adolescents exhibited a numerical trend in the same direction that did not reach significance (*t*(875) = 3.373, *p* = .051).

For excess path, both adults and adolescents showed lower excess path values on the first trial of Run 3 than Run 1 (Adults: *t*(864) = 4.591, *p* < .001; Adolescents: *t*(864) = 3.613, *p* = 0.02), although adolescents did not show corresponding accuracy-based signatures of prediction error or rapid updating. Together, these findings indicate that adults update and integrate spatial knowledge across runs, consistent with flexible, map-based representations. Adolescents show partial integration—evident in gradual improvements across runs and some decreases in excess path—but lack key signatures of immediate, map-based updating and integration. Children, by contrast, show little evidence of integration, despite performing above chance across changing task contingencies.

#### Within-run learning reveals distinct strategies across development

Finally, we examined within-run learning to test whether younger participants relied more on trial-by-trial feedback as task contingencies changed. When we tested this for all runs, children (*t*(875) = 1.714, *p* = .087) and adolescents (*p* = .089) exhibited a numerical trend toward within-run, across-trial improvements in accuracy on Run 2, whereas adults did not show this pattern (*p* > 0.143). Although these results did not reach significance, and were only observed for Run 2, they are consistent with a greater reliance on incremental, feedback-driven learning in younger participants.

When a new object set was introduced in Run 4, all age groups showed significant or trend-level evidence of within-run learning. Accuracy improved across trials for both adults (*t*(875) = 2.764, *p* = .005) and children (*t*(875) = 5.500, *p* < .001), with a similar but non-significant trend in adolescents (*t*(875) = 1.812, *p* = .07). Adults additionally showed decreases in excess path values (*t*(864) = 2.807, *p* = .005), whereas no such effects were observed on later runs (*p*s > .549), indicating that these trial-by-trial gains in adults were specific to the introduction of novel information.

Critically, despite these shared within-run effects, age groups differed in how learning accumulated across runs. As noted above, adolescents and adults showed greater overall improvements in scaled accuracy from Run 1 to Run 4 than children (significant or trend-level effects). To further characterize the pattern of trial-by-trial gains observed in Run 4, we conducted an exploratory analysis of navigation time. Relative to Run 1, adults showed a smaller decrease in navigation time than children in Run 4 (*B* = 2.249, SE = 1.018, *t* = 2.210, *p* = 0.028; **Figure S3, Panel B**), indicating that adults spent relatively longer navigating when confronted with the new object set. Thus, although adults showed within-run gains in accuracy, these gains were accompanied by relatively sustained navigation times, consistent with increased engagement. Taken together, these findings suggest that similar within-run accuracy gains may reflect different underlying strategies: adults appeared to actively integrate the new object set into an existing spatial framework, whereas children relied more on de novo, feedback-driven learning.

Taken together, these results point to a developmental shift from item-based learning in children, to partial integration in adolescence, to fully integrated, map-based representations in adulthood. Although adolescents exhibited gains in scaled accuracy and decreases in excess path values across runs, several markers point to an immature profile. These include the absence of prediction errors, lower overall accuracy and higher excess path values relative to adults, and comparable performance across proximal and distal conditions in scaled accuracy (**Figure S4, Panel A**) but not excess path (**Figure S4, Panel B**), indicating difficulty representing and navigating the global configuration. Moreover, relative to children, adolescents showed selective improvements in accuracy when new objects were introduced, conditions under which updating demands were reduced. This pattern is consistent with strong memory for specific prior experiences coupled with a still-developing ability to integrate across changing contingencies. Having established these behavioral differences in how spatial knowledge is acquired and expressed, we next examined whether these behaviors are associated with developmental differences in hippocampal temporal autocorrelation.

### Developmental increases in anterior hippocampal autocorrelation track distinct components of spatial knowledge

We examined whether differences in hippocampal autocorrelation tracked task performance, and whether this relationship varied by age. To derive brain-behavior associations, we modeled measures of performance (scaled accuracy and excess path) and autocorrelation at the run level, taking differences in navigation time and run-specific covariates into account in all models (see **Methods**). Based on the behavioral evidence above, we expected hippocampal autocorrelation to relate most strongly to performance when participants used prior spatial structure to guide behavior, a profile expressed most consistently in adults and more partially in adolescents. Scaled accuracy, which depends on retrieving object locations within the environment, was expected to draw on both hippocampal subregions in adults, reflecting posterior coding of precise spatial coordinates and anterior integration of those coordinates within a broader spatial structure. Excess path, which indexes efficient navigation through the global environment, was expected to rely more selectively on anterior hippocampus.

For scaled accuracy, a hierarchical mixed-effects model examined whether the association between temporal autocorrelation, performance, and hippocampal subregion varied across age groups. Importantly, the three-way age group x behavior x subregion interaction was not significant (*p*s > .088; **Table S11**), indicating that the relationship between performance and hippocampal subregion did not differ reliably across age groups. Nevertheless, the reduced model (**Table S12**) revealed several lower-order effects. Adults exhibited a stronger association between performance and temporal autocorrelation than adolescents (*B* = −0.023, SE = 0.010, *t* = −2.208, *p* = 0.030; **Table S12**) and children (*B* = −0.027, SE = 0.011, *t* = −2.462, *p* = 0.016; **Table S12**). There was also a performance x subregion interaction (*B* = 0.005, SE = 0.002, *t* = 3.160, *p* = 0.002; **Table S12**), indicating that the positive association between scaled accuracy and temporal autocorrelation was stronger in the anterior than posterior hippocampus. Finally, consistent with the prior age x subregion models, the relationship between temporal autocorrelation and age group continued to vary by hippocampal subregion even when accounting for behavior, with adults showing a larger anterior-versus-posterior difference than children (*B* = −0.734, SE = 0.075, *t* = −9.760, *p* < .001; **Table S12**) and adolescents (*B* = −0.544, SE = 0.065, *t* = −8.385, *p* < .001; **Table S12**). Notably, we replicated the same pattern of results when we used raw distance error, rather than scaled accuracy (**Table S13**).

Although the three-way interaction was not significant, the reduced model revealed significant age group x performance, subregion x performance, and age group x subregion interactions. We therefore conducted follow-up simple effects analyses to further characterize these lower-order effects, focusing on the association between scaled accuracy and temporal autocorrelation within each age group and hippocampal subregion. These analyses (**Figure 4**; **Table S14**) showed that, within anterior hippocampus, scaled accuracy positively tracked autocorrelation in adults (*B* = 0.029, SE = 0.010, *t* = 2.974, *p* = 0.004) and adolescents (*B* = 0.007, SE = 0.003, *t* = 2.142, *p* = 0.035), but not in children (*p* = 0.643). Conversely, within posterior hippocampus, scaled accuracy was associated with autocorrelation in adults (*B* = 0.025, SE = 0.010, *t* = 2.487, *p* = 0.015; **Table S14**), and not in the other age groups (*p*s > 0.548; **Table S14**).

**Figure 4.**
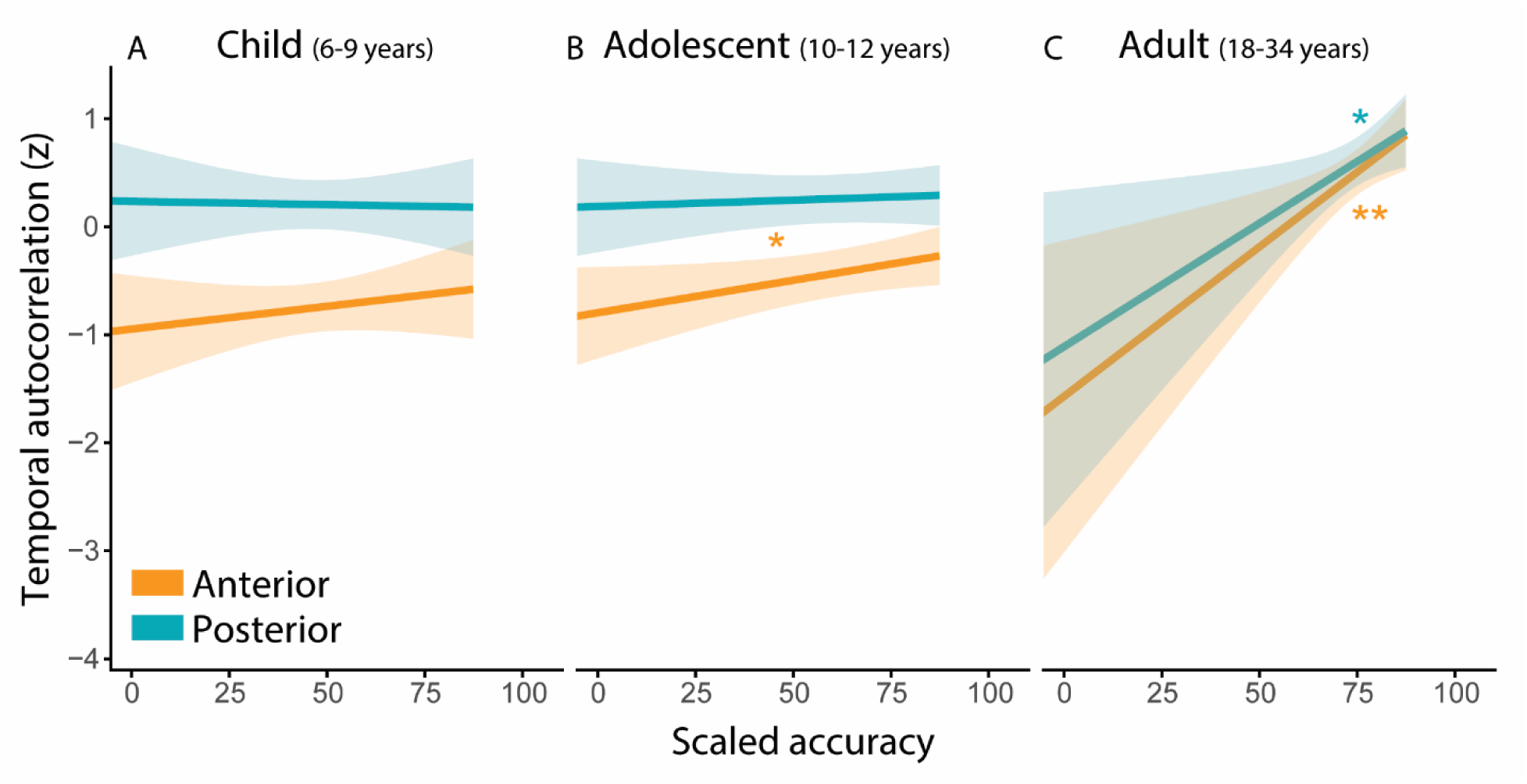
Follow-up simple-slopes analyses characterizing the associations between scaled accuracy and temporal autocorrelation, separately by age group and hippocampal subregion. Although the three-way performance x age group x subregion interaction was not significant, the analyses shown characterize the significant lower-order interactions, including: (1) the performance x age group interaction (stronger performance-autocorrelation associations in adults than in adolescents or children; *p*s < 0.030), (2) the performance x subregion interaction (stronger performance-autocorrelation associations in anterior than posterior hippocampus; ps < 0.030, and (3) the subregion x age group interaction (stronger age-related increases in anterior relative to posterior hippocampal temporal autocorrelation). **A)** In children, temporal autocorrelation was not related to scaled accuracy in either subregion (*p*s > 0.616). **B**) In adolescents, scaled accuracy was positively associated with temporal autocorrelation in anterior (*B* = 0.007, SE = 0.003, *t* = 2.142, *p* = 0.035) but not posterior hippocampus (*p* = 0.548). **C**) In adults, scaled accuracy was positively associated with temporal autocorrelation in both hippocampal subregions (anterior: *B* = 0.029, SE = 0.010, *t* = 2.974, *p* = 0.004; posterior: *B* = 0.025, SE = 0.010, *t* = 2.487, *p* = 0.015). Lines and 95% CI’s derived from hierarchical mixed-effects models with age group, scaled accuracy, and all covariates as predictors. \**p* < .05, \*\**p* < .01. Each participant provided four observations (anterior and posterior hippocampus in left and right hemisphere), resulting in 332 measurements contributing to the model.

To test whether hippocampal autocorrelation was specifically related to across-run integration or to local feedback-based improvement, we next examined within-run changes in scaled accuracy. Within-run gains provide a more limited index of spatial learning because they can reflect trial-by-trial correction based on recent feedback, rather than stabilization or generalization of spatial structure across runs, which we hypothesized in children. We found that within-run accuracy gains in children and adolescents were not associated with temporal autocorrelation (*p* = 0.347; **Table S15**). Thus, hippocampal autocorrelation did not appear to index local, feedback-based improvement within a run, but rather the more stable spatial representations that supported performance across changing task contexts.

For excess path, which indexes efficient navigation across the broader spatial environment, we observed a three-way interaction between age group, subregion, and excess path in predicting temporal autocorrelation (Child: *B* = 0.050, SE = 0.011, *t* = 4.661, *p* < .001; Adolescent: *B* = 0.059, SE = 0.011, *t* = 5.459, *p* < .001; **Table S16**). Follow-up analyses revealed that, in adults, greater anterior—but not posterior— hippocampal autocorrelation was associated with more efficient navigation (*B* = −0.072, SE = 0.018, *t* = −3.918, *p* < .001; **Figure 5C**; **Table S17**), consistent with their use of a fully integrated map to traverse more direct paths to their goal. In contrast, adolescents (*p*s > .088) and children (*p*s > .486) showed no relationship between autocorrelation and excess path (**Figure 5A-B**; **Table S17**), paralleling behavioral evidence that only adults exhibited consistently efficient navigation across proximal and distal conditions. Together, these findings suggest that hippocampal autocorrelation becomes increasingly linked to scaled accuracy across development, with mature performance in adults supported by both anterior and posterior hippocampal contributions. In contrast, efficient navigation through the broader spatial environment (excess path) showed an adult-specific association with anterior hippocampal autocorrelation, suggesting that the role of anterior hippocampus in supporting globally integrated, map-based navigation emerges later in development.

**Figure 5.**
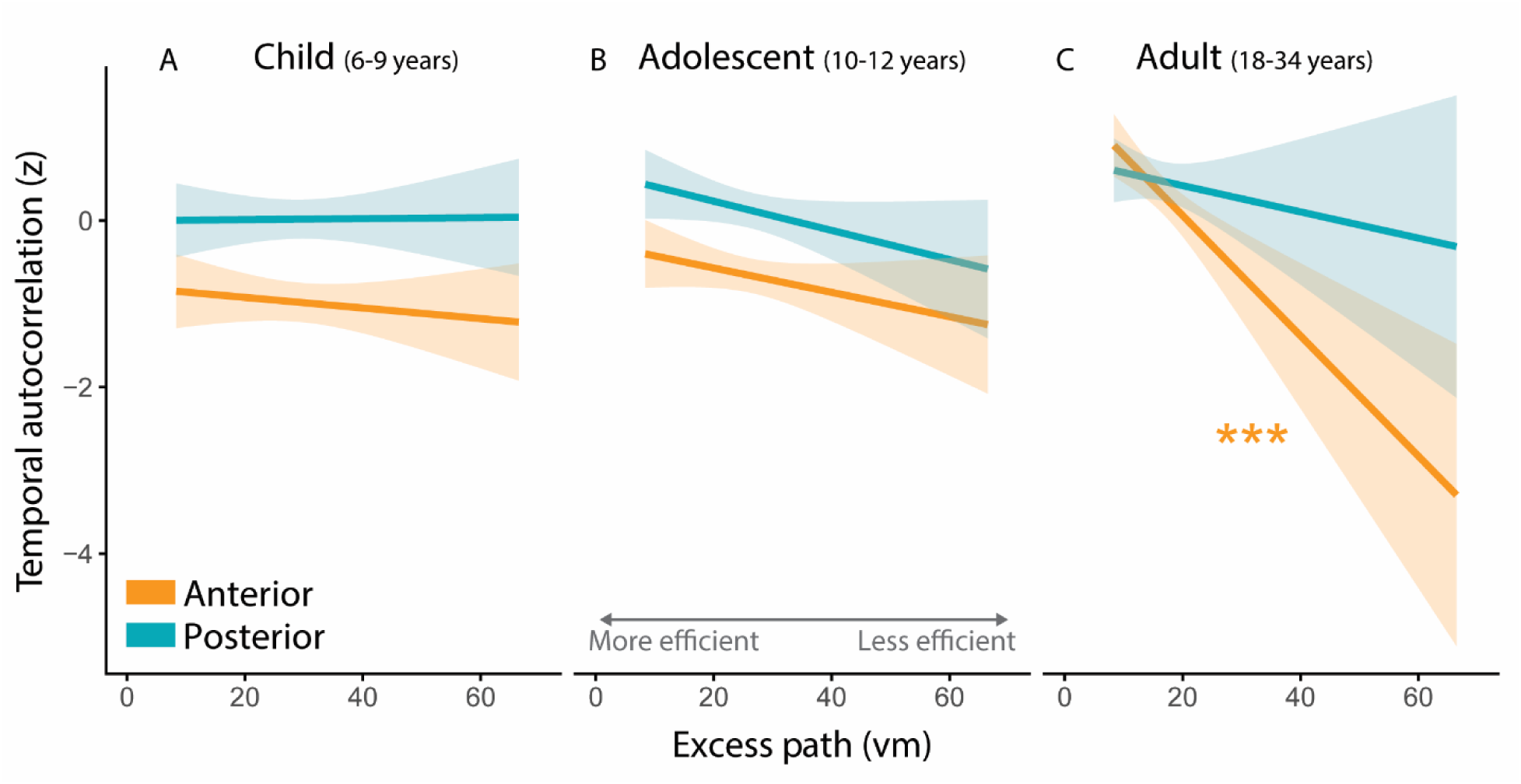
Follow-up simple-slopes analyses characterizing the significant excess path x age group x subregion interaction, with associations between excess path and temporal autocorrelation plotted separately by age group and hippocampal subregion. **A**) Temporal autocorrelation does not correspond to excess path in either hippocampal subregion in children (*p*s > 0.486) nor **B**) adolescents (*p*s > 0.088). **C**) In adults, greater anterior hippocampal temporal autocorrelation relates to lower excess path (*B =* −0.072, SE = 0.018, *t* = −3.918, *p* < .001) while posterior hippocampal autocorrelation was not associated with performance (*p* = 0.395). Lines and 95% CI’s are derived from mixed-effects models with age group, excess path, and all covariates as predictors. One participant was excluded from all excess path analyses due to missing data. Each participant provided four observations (anterior and posterior hippocampus in left and right hemisphere), resulting in 328 measurements represented above.

## Discussion

Successful navigation requires both learning individual locations and recognizing stable spatial structure across changing experiences. A central question is how children and adolescents acquire this capacity to integrate spatial knowledge into representations that can be updated and generalized across changing environments. Here, we show that this developmental transition is associated with complementary changes in anterior and posterior hippocampal function. Across two independent samples, posterior hippocampal temporal autocorrelation remained largely stable with age, whereas anterior hippocampal autocorrelation increased into adulthood. Yet the behavioral relevance of these signals differed across development: adults showed parallel anterior and posterior hippocampal associations with precise object-location memory, while anterior hippocampal autocorrelation uniquely predicted efficient navigation trajectories. These findings suggest that the later emergence of anterior hippocampal computations supports navigation of novel routes through a changing landscape to remembered locations.

Interpreting these hippocampal signals required a task that distinguished remembering locations from using an integrated map. Prior work shows that children and adolescents increasingly shift from route-based or local spatial strategies toward more flexible representations that support shortcuts, path integration, and cognitive maps (Brucato et al., 2022; Burles et al., 2020; Nazareth et al., 2018). Our task extends this work by dissociating scaled accuracy, which indexes memory for object locations, from excess path, which indexes whether participants used broader spatial structure to plan efficient routes from novel starting positions. This distinction proved critical. All age groups performed above chance, showing that even children could learn object locations as task contingencies changed.

However, above-chance performance did not imply a common representational strategy. Children showed some within-run gains consistent with feedback-driven learning, but little evidence of across-run updating or generalization. Adolescents showed partial use of prior spatial structure, with across-run gains and more adult-like scaled accuracy across proximal and distal objects, but without clear evidence of rapid structure-based updating or efficient navigation across conditions. Adults showed the clearest evidence of map-based behavior, combining precise object-location memory with efficient trajectories and rapid use of prior spatial structure. Thus, developmental differences were evident in not only how accurately participants remembered locations, but whether those locations were embedded in a flexible spatial representation that could guide navigation as the environment changed.

Temporal autocorrelation allowed us to ask which hippocampal computation supported the shift toward map-based behavior. Anterior hippocampal structure, activation, and connectivity mature later than posterior hippocampus, consistent with the prolonged emergence of memory integration and flexible inference (Calabro et al., 2020; Demaster et al., 2016; DeMaster & Ghetti, 2013; Schlichting et al., 2016; Tang et al., 2020; Xie et al., 2024). However, these measures do not specify how hippocampal activity becomes organized over the temporal unfolding of experience to support stable representations that link distinct events. Temporal autocorrelation provides such a window by indexing the persistence of neural activity patterns over time, a property thought to support integration across events (Paz et al., 2010; Rait et al., 2025; Raut et al., 2020). To capture this persistence across navigation trials, we measured autocorrelation over full task runs. Greater run-level autocorrelation is therefore consistent with more stable neural activity across distinct experiences, potentially supporting the global representation of all experience, not just a single event.

Across both resting-state and task-based fMRI samples, anterior hippocampal autocorrelation increased with age, identifying a computational signature of prolonged anterior hippocampal maturation. The replication across rest and task suggests that autocorrelation captures a stable feature of hippocampal maturation, consistent with prior evidence that temporal autocorrelation is reliable across repeated assessments in adults (Bouffard et al., 2023). Notably, this developmental increase was not observed in posterior hippocampus, indicating that maturation was not a global feature of hippocampal autocorrelation. Furthermore, while posterior hippocampal autocorrelation appeared relatively stable across age, its behavioral relevance emerged only in adulthood, underscoring the need to distinguish between developmental change in a neural signal from how that signal supports behavior.

The observation that posterior hippocampal autocorrelation was associated with scaled accuracy only in adults provides important insight into how precise spatial memory becomes behaviorally useful. Prior developmental work shows that posterior hippocampal representations become increasingly precise and memory-relevant during development (Callaghan et al., 2021). Our findings extend this work by showing that, although the similarity of posterior hippocampal activity patterns across time remained stable with age, its behavioral relevance changed across development. Posterior autocorrelation tracked scaled accuracy only in adults, indicating that an age-invariant posterior signal is not sufficient, on its own, to support adult-like spatial precision. Rather, posterior coding may become most behaviorally useful when later-maturing anterior hippocampal computations that integrate individual object locations within broader spatial structure are also available. This joint anterior-posterior association is notable because precise object-location memory had to be maintained across changes in egocentric viewpoint, distal cue orientation, and object set. Thus, mature scaled accuracy appears to depend on complementary long-axis contributions: posterior hippocampus anchors precise object-location information, whereas anterior hippocampus embeds those locations within a flexible map that can be updated across changing contexts (Evensmoen et al., 2015; Nadel et al., 2013; Poppenk et al., 2013).

If scaled accuracy reflected maintenance of precise object-location memories within a changing spatial structure, excess path provided a more stringent test of whether participants used that structure to guide behavior, by locating themselves within the broader environment and planning direct trajectories to remembered goals. Only adults showed an association between lower excess path and greater anterior hippocampal autocorrelation. This anterior-specific relationship extends prior adult work showing greater anterior than posterior hippocampal autocorrelation during demanding navigation (Brunec et al., 2018) by directly linking this temporal coding property to efficient behavior. It also converges with evidence that slower hippocampal autocorrelation supports how temporal memories are organized in adults (Rait et al., 2025), suggesting that temporally stable anterior hippocampal activity may organize experiences across space as well as time. Thus, mature anterior hippocampal autocorrelation appears to support not only memory for where objects are, but the flexible use of those memories to navigate efficient routes through a changing environment, consistent with maintenance of an integrated representation.

Our results in children help further clarify what temporal autocorrelation does— and does not—index in this task. Children performed above chance and showed initial within-run gains, indicating that they could learn object locations from feedback. However, they showed little evidence of across-run updating or generalization, and neither anterior nor posterior hippocampal autocorrelation was reliably related to behavior. Thus, hippocampal autocorrelation does not appear to be a generic marker of above-chance learning. Instead, it appears to track spatial representations that are stable enough to guide behavior beyond the immediate context in which learning occurred. This interpretation aligns with evidence that younger learners may preserve distinct experiences when relational structure is uncertain, with flexible integration emerging gradually as hippocampal and cortical systems mature (Benear et al., 2022; Keresztes et al., 2018; Schlichting et al., 2022; Varga et al., 2025). However, while these null effects are consistent with developmental change, they should be interpreted cautiously, as the present data do not establish that the association between hippocampal coding and behavior is absent in children more generally, but only that we did not detect it in the current approach. Future work is needed to determine what alternative neural mechanisms support navigation in children.

Adolescents showed a more advanced but still immature profile. Unlike children, adolescents showed across-run gains, suggesting use of prior spatial structure. They also showed an association between anterior hippocampal autocorrelation and scaled accuracy, indicating that emerging anterior stability supported memory for object locations within the changing environment. However, adolescents did not show the full adult profile: they lacked robust evidence of rapid structure-based updating, parallel anterior-posterior associations with scaled accuracy, and anterior associations with excess path. This pattern helps reconcile the present findings with prior work showing that early adolescents can approach adult-like navigation performance in stable virtual environments (Brucato et al., 2022; Nazareth et al., 2018). Early adolescence may support cognitive maps under relatively stable conditions, but adult-like flexibility is still developing when spatial knowledge must be updated and generalized more broadly (Friend et al., 2026).

Taken together, these findings provide a neural account of a longstanding developmental idea: the growing ability to integrate information across time and space transforms how individuals represent and use knowledge across development (Brucato et al., 2022; Friend et al., 2025; Nazareth et al., 2018; Piaget, 1970; Varga et al., 2019; Varga & Bauer, 2017). By linking hippocampal temporal autocorrelation to dissociable behavioral indices of object-location memory and efficient route planning, the present study provides a developmental test of theories proposing functional specialization along the hippocampal long axis (Moser & Moser, 1998; Poppenk et al., 2013; Strange et al., 2014). Although posterior hippocampal autocorrelation remained largely stable with age, mature spatial behavior was linked to parallel anterior-posterior associations with object-location memory and a uniquely anterior association with efficient navigation in adulthood. Thus, flexible memory development reflects more than increasingly accurate recall of individual locations; it reflects the emergence of hippocampal representations that remain stable across changing experiences, allowing prior spatial knowledge to support novel routes, updating, and generalization as the world changes.

## Conflict of Interests Statement

The authors declare no competing interests.

## Acknowledgments

The authors extend their gratitude to Katharine Guarino and Kim Nguyen for assistance with piloting and data collection. In addition, GPT-5.1 was used by the first author (NLV) for AI-assisted copy editing of the final main text, exclusively for improving sentence structure and readability and not for generative editorial work. Finally, we thank the families and participants.

## Data and Code Accessibility

The data and custom code that support the analyses of this study are publicly available on the Open Science Framework (https://osf.io/g7qne).

## Author Contributions

[H.E.R., M.L.S., K.R.S., C.A.C. and A.R.P.] designed research. [H.E.R., K.R.S. and M.L.S.] performed research. [O.W.F., N.L.V., and A.M.D.] analyzed data. [N.L.V. and O.W.F.] drafted the paper, and [A.R.P.] revised the manuscript for review and finalization by all other authors.

## Funding Information

This work was supported by funding from NIMH R01 grant MH100121 (ARP) and National Science Foundation Division of Graduate Education grant DGE2137420 (OWF). The funders had no role in study design, data collection and analysis, or preparation of the manuscript.

## Supplemental Materials

### Scaled accuracy calculation

We first quantified the raw distance error in virtual meters (vm) between the participant’s recalled location and the target object’s actual location for each trial (**Figure 1C**). We then transformed these raw error scores into a scaled score to account for the non-uniform chance error that arises from the circular nature of the virtual environment, wherein, even if responses were completely random, targets located near the center would, on average, produce smaller distance errors than targets located near the boundary. Thus, because two target objects within the navigation task were always located near the boundary of the arena (i.e., distal objects) and two were always located near the center (i.e., proximal objects), participants would demonstrate improved accuracy for proximal objects relative to distal even if navigating completely randomly (**Figure 1A**). As such, we computed and report scaled accuracy scores that adjust for the possible deviation that could be observed for each location.

Specifically, following prior work (Bullens et al., 2010), we discretized the circular arena into 88 possible response locations arranged in a grid (i.e., equidistant from each other). For each target location, we then iteratively computed the Euclidean distance between that target location and each of the 88 possible response locations (**Figure S2, Panel B**; left arena), averaging those distances to derive a single value for how much distance error should be expected from that target location based on random chance. Next, for each behavioral trial for each participant, we translated the error into a scaled accuracy score, defined as 100*(chance error – observed error)/chance error, resulting in a single scaled accuracy value for each trial, with 0 representing random chance and 100 representing perfect navigation. To confirm that discretizing the arena into response locations did not bias results based on grid granularity, we repeated the analysis using a finer grid with twice as many locations (**Figure S2, Panel B**; right arena). This up-sampling yielded identical results, so all subsequent analyses were conducted using the original 88-location grid. Finally, we also repeated all analyses using the raw distance error scores, and the overall pattern of results remained unchanged (**Table S13**).

### Excess path calculation

Excess path was calculated for each trial, quantified as the difference between a Euclidean path between the spawn and target locations (a maximally efficient path) and the total distance a participant travelled from spawn to target, which summed navigation and feedback (in vm). As such, lower values reflect more efficient navigation, while higher values reflect less efficient navigation. Importantly, excess path values controlled for differences in object location accuracy because the total distance traveled in each trial included the corrective feedback they received after indicating their recalled location. In other words, trials in which participants traversed short distances, but did not accurately recall the location, were not artificially measured as highly efficient.

Excess path distance between traversed and optimal paths was used here instead of a ratio between the traversed and optimal path length because ratio measures can become disproportionately skewed when optimal paths are short. Although the distance between traversed and optimal paths could also inflate differences at the level of individual trials, the randomized spawn locations of our design ensured that the total optimal path distance within each run remained comparable across participants (mean optimal distance = 39.501, SD = 2.312). Accordingly, the absolute difference metric was better suited to our run-by-run modeling approach. That said, to further account for any remaining variance in trial geometry, the average optimal path distance across each run was included as a covariate in our primary analyses and did not significantly explain variance in performance (p = 0.565).

### Hippocampal autocorrelation across varying temporal lags

Our primary results highlight developmental differences in hippocampal temporal autocorrelation, assessed by iteratively computing the correlation of neural activity at successive TRs (i.e., across a lag of one TR). In addition, we assessed the extent to which age differences in hippocampal autocorrelation corresponded to the temporal lag across which autocorrelation was measured. To that end, we calculated temporal autocorrelation for each run across lags of 2, 3, 4, and 10 TRs (i.e., 4-20 seconds) with an identical approach to our primary lag-1 analyses.

Within the resting-state sample, lag did not interact with age or subregion via two-way or three-way interactions (ps > 0.291). Moreover, we identified identical age x subregion interactions to our primary lag-1 analyses (anterior hippocampal autocorrelation increasing with age more than posterior hippocampus) at every lag: lag 2 (B = 0.084, SE = 0.006, t = 13.761, p < .001), lag 3 (B = 0.085, SE = 0.006, t = 13.769, p < .001), lag 4 (B = 0.084, SE = 0.006, t = 13.702, p < .001), and lag 10 (B = 0.084, SE = 0.006, t = 13.654, p < .001).

Within the task-based sample, when the developmental group was examined continuously (6-12 years), lag similarly did not interact with age in years and months or subregion (ps > 0.424). We identified the same age x subregion interaction with anterior hippocampal autocorrelation increasing more with age than posterior autocorrelation at lag 2 (B = 0.079, SE = 0.019, t = 4.286, p < .001), lag 3 (B = 0.080, SE = 0.018, t = 4.341, p < .001), lag 4 (B = 0.080, SE = 0.018, t = 4.312, p < .001), and lag 10 (B = 0.79, SE = 0.018, t = 4.262, p < .001).

Lag similarly did not interact with subregion or age when age was considered as a categorical predictor (ps > 0.087). Again, results were identical to our primary results at all lags, with age group x subregion interactions suggesting greater anterior autocorrelation with age: lag 2 (child: B = −0.875, SE = 0.060, t = −14.426, p < .001; adolescent: B = −0.606, SE = 0.061, t = −9.906, p < .001), lag 3 (child: B = −0.884, SE = 0.061, t = −14.571, p < .001; adolescent: B = −0.615, SE = 0.061, t = −10.067, p < .001), lag 4 (child: B = −0.882, SE = 0.060, t = −14.573, p < .001; adolescent: B = −0.614, SE = 0.060, t = −10.075, p < .001), and lag 10 (child: B = −0.879, SE = 0.060, t = −14.521, p < .001; adolescent: B = −0.614, SE = 0.060, t = −10.077, p < .001).

**Figure S1.**
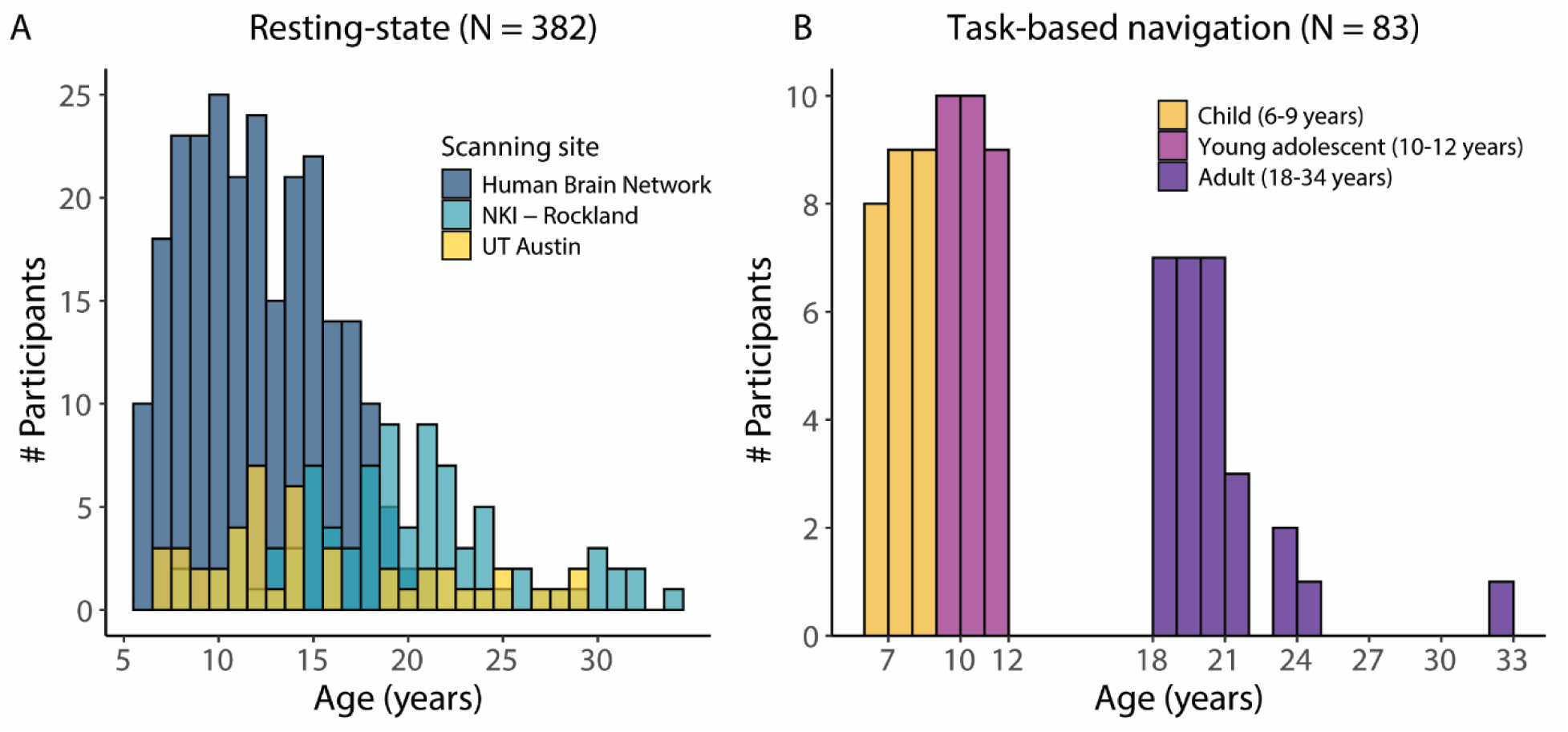
Age and site distributions for the final participant samples. Histograms are plotted separately for the **A)** resting-state and **B)** task-based navigation samples. See Materials and Methods (Participants section) for extended description of samples and inclusion criteria.

**Figure S2.**
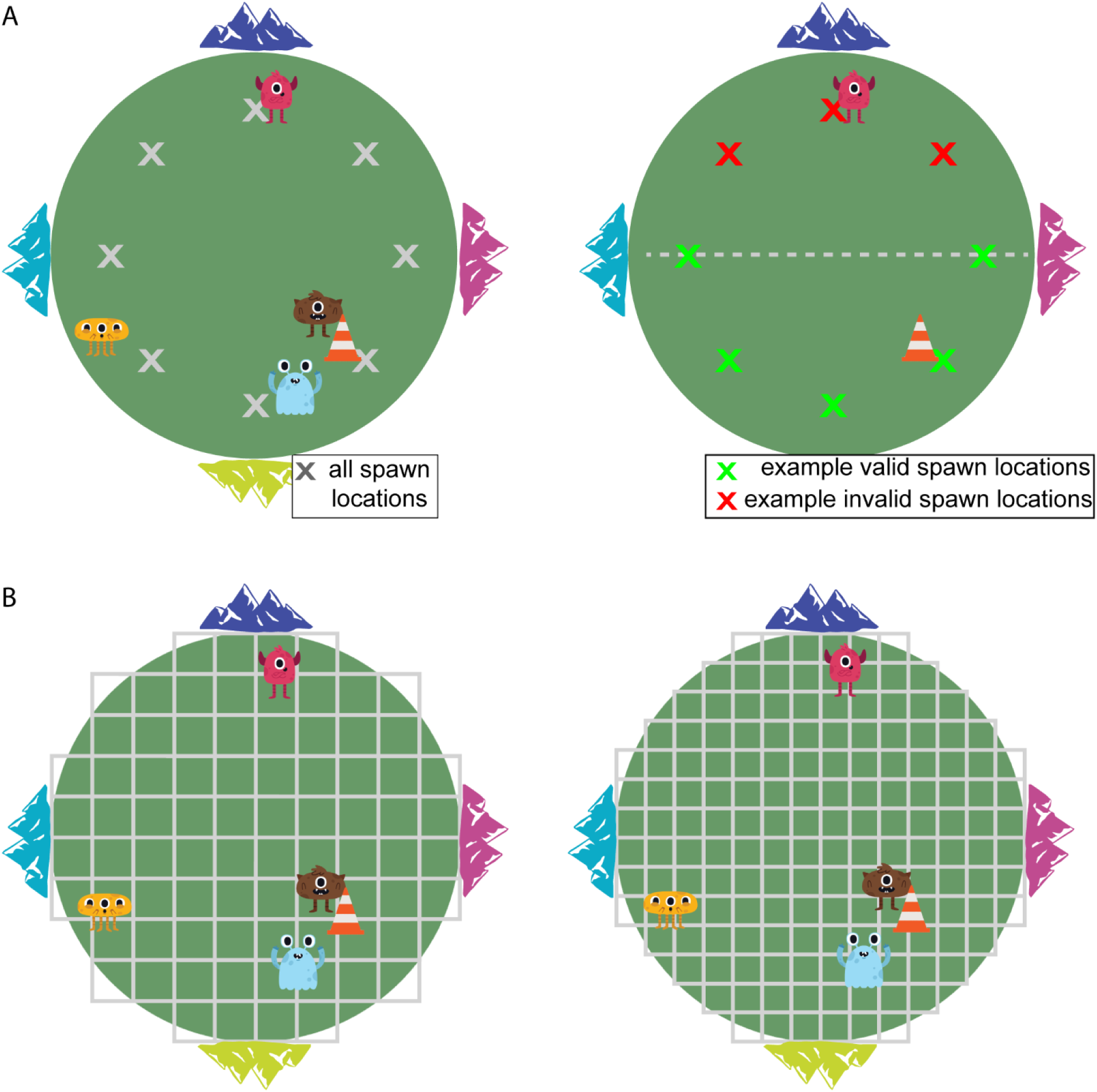
**A) Possible spawn locations.** Spawn locations included eight possible locations equally spaced about the arena (left arena - grey x’s). For each trial, one of five possible spawn locations within the opposite half of the arena from the target object (i.e., valid spawn locations: right arena, green x’s) was pseudo-randomly selected as the spawn location. As such, participants were never spawned in direct proximity to a target location. Moreover, the distance and angle between the spawn and target location differed between each trial. Critically, participants were never spawned in the same location for the same target object within a run (three different spawn locations per object within each run). This constraint required navigation along novel paths on every trial. **B) Possible response locations used to calculate scaled accuracy.** Depiction of the discretized arena used to calculate chance accuracy. For each square in the grid, distance was calculated between the center of the square and the target location. These values were averaged to compute the distance error that would be expected by random chance. The left arena demonstrates discretization of spatial arena using 88 equally sized squares (primary analyses), while the right arena demonstrates higher-resolution discretization with 176 smaller squares. Doubling the number of squares in the grid did not affect results (see above; **Scaled accuracy calculation**).

**Figure S3.**
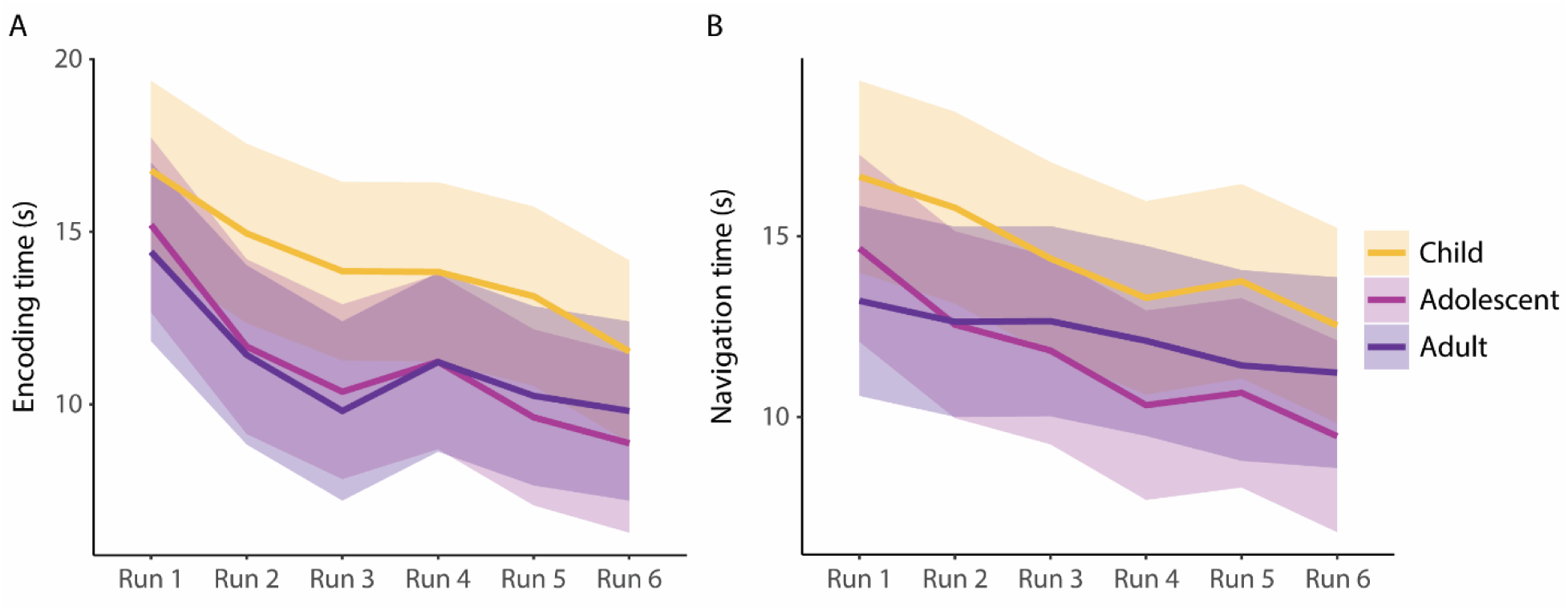
**A) Object encoding time during learning trials was consistent across age groups.** While we primarily assessed behavior during the test trials of the navigation task, the first four learning trials of each run allowed participants to spend as much time as needed navigating to the visually observable object (Figure 1A). Thus, age-related differences in time spent initially encoding the object locations could have emerged during these trials, which would point to potential differences in motor abilities or exploratory tendencies. To test for differences in encoding time by age group and run, we conducted mixed-effects models with average encoding time per run as the outcome variable and both main effect and interaction terms for age group, run, and condition (proximal versus distal) as predictors, in addition to a participant-level random intercept. Importantly, navigation time during encoding trials did not significantly differ by age group (ps > 0.64), suggesting that participants of all ages received equal exposure to target locations during learning trials. For all age groups, time spent navigating during encoding trials decreased across the duration of the experiment (i.e., from Runs 1-6; B = −5.232, SE = 1.185, t = −4.415, p < .001). Age group did not interact with run or condition (ps > 0.08), suggesting all groups received comparable exposure to learning trials across the duration of the experiment. **B) Differences in navigation time during test trials by age group.** Age groups did not significantly differ in navigation time during test trials (finding + feedback; Fig. 1B). All age groups spent progressively less time navigating during test trials across the task (Run 6 - Run 1: B = −4.116, SE = 0.768, t = −5.361, p < .001). We also identified two run x age group interactions indicating that, relative to Run 1, adults’ navigation time decreased less than children during Run 4 (B = 2.249, SE = 1.018, t = 2.210, p = 0.028) and during Run 6 (B = 2.111, SE = 1.039, t = 2.033, p = 0.043). Because of these age differences, number of TRs was included as a control measure in all models assessing age differences in temporal autocorrelation (see **Tables S5-S17**) but was not associated with temporal autocorrelation in any model.

**Figure S4.**
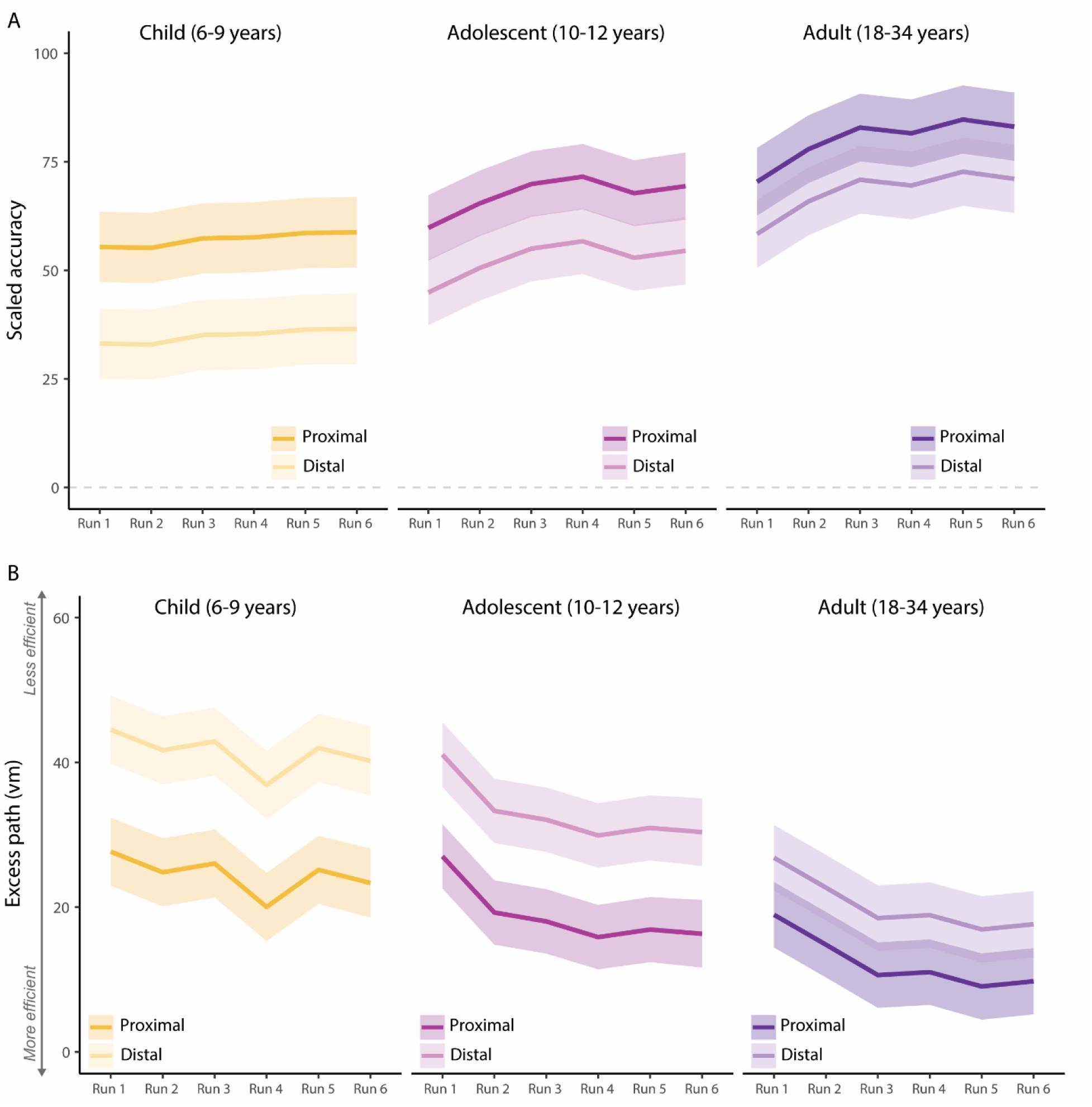
**A) Scaled accuracy differences based on condition and age group.** To assess differences in scaled accuracy on proximal versus distal trials, we implemented mixed-effects models with scaled accuracy as the outcome variable and both main effect and interaction terms for age group, run, and condition (proximal versus distal) as predictors. We found a main effect of condition such that all age groups were more accurate within the proximal condition than distal condition (B = 22.253, SE = 1.916, t = 11.618, p = < .001). We also identified age x condition interactions indicating that adult and adolescent performance differed less between the two conditions than children’s performance (adult: B = −10.236, SE = 2.664, t = −3.843, p < .001; adolescent: B = −7.374, SE = 2.603, t = −2.803, p = .005). Age group differences when accounting for condition were identical to those observed when collapsing across conditions (Figure 3A), with adults more accurate than children (t(81) = 5.964, p < .001) and adolescents in both conditions (t(81) = 3.130, p = .007), and adolescents more accurate than children in both conditions (t(81) = 2.990, p = 0.011). We did not identify any three-way interactions between age group, condition, and run. Finally, we conducted confirmatory analyses to ensure that each age group navigated more accurately than chance for both proximal and distal trials within each run. To that end, we conducted one-sample t-tests for scaled accuracy within each age group, run, and condition combination. All age groups were more accurate than chance on all six runs within both proximal and distal conditions (ps < .001). **B) Excess path differences based on condition and age group.** We implemented the same analysis to examine condition effects of excess path. We found a main effect of condition such that all age groups showed lower excess path values within proximal trials (B = −16.859, SE = 1.458, t = −11.560, p < .001), consistent with their increased scaled accuracy for proximal trials. We also identified an age x condition interaction indicating that adults’ excess path values differed less between the two conditions than children’s (adult: B = 8.977, SE = 2.019, t = 4.427, p < .001), while adolescents did not differ from either age group (ps > 0.164). Again, age group differences when accounting for condition were identical to those observed when collapsing across conditions (Figure 3B), with adults showing lower excess path values than children (t(80) = 7.524, p < .001) and adolescents in both conditions (t(80) = 4.465 p = .007), and adolescents showing lower excess path values than children in both conditions (t(81) = 3.196, p = 0.006).

**Table S1.** Image acquisition parameters by scanning site.

|  | HBN (Rest) | NKI-RS (Rest) | UT Austin (Rest) | UT Austin (Task) |
| --- | --- | --- | --- | --- |
| TR (s) – T1w Anatomical | 2.5 | 2.5 | 1.9 | 1.9 |
| TE (ms) – T1w Anatomical | 3.15 | 3.5 | 2.43 | 2.43 |
| Voxel Size (mm) – T1w Anatomical | 0.8x0.8x0.8 | 1.0x1.0x1.0 | 1.0x1.0x1.0 | 1.0x1.0x1.0 |
| Flip Angle (deg) – T1w Anatomical | 8 | 8 | 9 | 9 |
| FoV (mm) – T1w Anatomical | 256 | 256 | 256 | 256 |
| Runs – T2*w Functional | 2 | 1 | 2 | 6 |
| TRs (#) – T2*w Functional | 375 | 404 | 160 | 220 |
| TR (s) – T2*w Functional | 0.8 | 1.4 | 1.89 | 2 |
| TE (ms) – T2*w Functional | 30 | 30 | 30 | 30 |
| Voxel Size (mm) – T2*w Functional | 2.43x2.43x2.40 | 2.0x2.0x2.0 | 1.5x1.5x1.5 | 2.0x2.0x2.0 |
| Flip Angle (deg) – T2*w Functional | 31 | 65 | 60 | 71 |
| FoV (mm) – T2*w Functional | 204 | 224 | 220 | 220 |
| Multiband Acceleration Factor – T2*w Functional | 6 | 4 | 4 | 3 |
| GRAPPA Factor – T2*w Functional | NA | NA | 2 | 2 |
| TR (ms) – B0 Fieldmap | 5.301 | NA | 0.589 | 0.706 |
| TE (ms) – B0 Fieldmap | 51.2 | NA | 5 | 5.0/7.46 |
| Voxel size (mm) – B0 Fieldmap | 2.4x2.4x2.4 | NA | 1.5x1.5x2.0 | 1.7x1.7x2.0 |
| Flip angle (deg) – B0 Fieldmap | 90 | NA | 5 | 5 |
| FoV (mm) – B0 Fieldmap | 202 | NA | 192 | 220 |

*Note.* NKI-RS scans did not include fieldmap images.

**Table S2.** Continuous age differences in temporal autocorrelation by subregion during rest.

| Fixed effects | Coef. Estimate | Standard Error | t value | p-value |
| --- | --- | --- | --- | --- |
| Intercept | 1.699 | 0.691 | 2.457 | 0.014* |
| Age (years and months) | -0.007 | 0.008 | -0.863 | 0.388 |
| Subregion (anterior) | -2.228 | 0.139 | -16.009 | <.001*** |
| Age-Subregion (anterior) interaction | 0.079 | 0.006 | 12.774 | <.001*** |
| Hemisphere (right) | 0.073 | 0.033 | 2.217 | 0.027* |
| tSNR | 0.001 | 0.005 | 0.276 | 0.783 |
| Number TRs | 0.000 | 0.001 | -0.059 | 0.953 |
| Number voxels | 0.002 | 0.000 | 4.082 | <.001*** |
| Avg. voxel distance – x direction | 0.205 | 0.130 | 1.581 | 0.114 |
| Avg. voxel distance – y direction | -0.646 | 0.085 | -7.605 | <.001*** |
| Avg. voxel distance – z direction | -0.365 | 0.080 | -4.546 | <.001*** |
| Mean framewise displacement | -0.374 | 0.686 | -0.545 | 0.586 |
| Mean absolute translation | 0.404 | 0.470 | 0.860 | 0.390 |
| Scanning site | -0.123 | 0.137 | -0.900 | 0.368 |

*Note:* \*p < .05. \*\*p < .01. \*\*\*p < .001. Model specification: temporal\_autocorrelation ~ intercept + age + subregion + age:subregion + hemisphere + tSNR + number\_TRs + number\_voxels + dist\_x + dist\_y + dist\_z + mean\_fd + mean\_abs\_trans + site + (1|subject/run)

**Table S3.** Subregion-specific follow-up analysis of age differences in temporal autocorrelation during rest.

| Effect | Coef. Estimate | Standard Error | t value | p-value |
| --- | --- | --- | --- | --- |
| Age (years and months) - Anterior | 0.072 | 0.008 | 8.857 | <.001*** |
| Age (years and months) - Posterior | -0.007 | 0.008 | -0.863 | 0.388 |

*Note:* \*p < .05. \*\*p < .01. \*\*\*p < .001. Slopes extracted directly from aggregate model (Table S2) using `sim_slopes` function of `interactions` package in R.

**Table S4.** Control analysis of temporal autocorrelation in parahippocampal cortex in resting-state sample.

| Fixed effects | Coef.<br>Estimate | Standard<br>Error | t value | p-value |
| --- | --- | --- | --- | --- |
| Intercept | -1.066 | 0.760 | -1.404 | 0.161 |
| Age (years and months; Rest) | -0.001 | 0.007 | -0.087 | 0.930 |
| Hemisphere (right) | -0.126 | 0.022 | -5.679 | < .001 |
| tSNR | 0.010 | 0.004 | 2.687 | 0.007** |
| Number TRs | 0.000 | 0.002 | 0.220 | 0.826 |
| Number voxels | -0.002 | 0.001 | -3.062 | 0.002*** |
| Avg. voxel distance – x direction | 0.586 | 0.119 | 4.920 | < .001*** |
| Avg. voxel distance – y direction | 0.154 | 0.088 | 1.743 | 0.082 |
| Avg. voxel distance – z direction | -0.078 | 0.093 | -0.842 | 0.400 |
| Mean framewise displacement | 0.161 | 0.066 | 2.447 | 0.015* |
| Mean absolute translation | 0.038 | 0.086 | 0.445 | 0.657 |
| Site (UT Austin) | -0.162 | 0.131 | -1.236 | 0.217 |
| Site (NKI-RS) | -0.038 | 0.493 | -0.077 | 0.939 |

*Note:* \*p < .05. \*\*p < .01. \*\*\*p < .001. Model specification: temporal\_autocorrelation ~ intercept + age + + hemisphere + tSNR + number\_TRs + number\_voxels + dist\_x + dist\_y + dist\_z + mean\_fd + mean\_abs\_trans + (1|subject/run)

**Table S5.** Continuous age differences in temporal autocorrelation by subregion during navigation task within developmental group.

| Fixed effects | Coef. Estimate | Standard Error | t value | p-value |
| --- | --- | --- | --- | --- |
| (Intercept) | -2.450 | 0.714 | -3.433 | < .001*** |
| Age (years and months) | 0.039 | 0.033 | 1.186 | 0.240 |
| Subregion (anterior) | -1.940 | 0.245 | -7.903 | < .001*** |
| Age-Subregion interaction | 0.079 | 0.018 | 4.322 | < .001 |
| Hemisphere (right) | 0.171 | 0.031 | 5.477 | < .001*** |
| tSNR | 0.054 | 0.002 | 22.850 | < .001*** |
| Number TRs | 0.000 | 0.001 | -0.346 | 0.729 |
| Number voxels | 0.002 | 0.001 | 2.070 | 0.039* |
| Avg. voxel distance – x direction | 0.182 | 0.166 | 1.097 | 0.273 |
| Avg. voxel distance – y direction | -0.419 | 0.113 | -3.719 | < .001*** |
| Avg. voxel distance – z direction | -0.048 | 0.107 | -0.446 | 0.656 |
| Mean framewise displacement | 2.070 | 1.010 | 2.047 | 0.042* |
| Mean absolute translation | -1.900 | 0.859 | -2.213 | 0.028* |
| Run configuration | 0.158 | 0.119 | 1.324 | 0.187 |

*Note:* \*p < .05. \*\*p < .01. \*\*\*p < .001. Model specification: temporal\_autocorrelation ~ intercept + age + subregion + age:subregion + hemisphere + tSNR + number\_TRs + number\_voxels + dist\_x + dist\_y + dist\_z + mean\_fd + mean\_abs\_trans + run\_configuration + (1|subject/run)

**Table S6.**
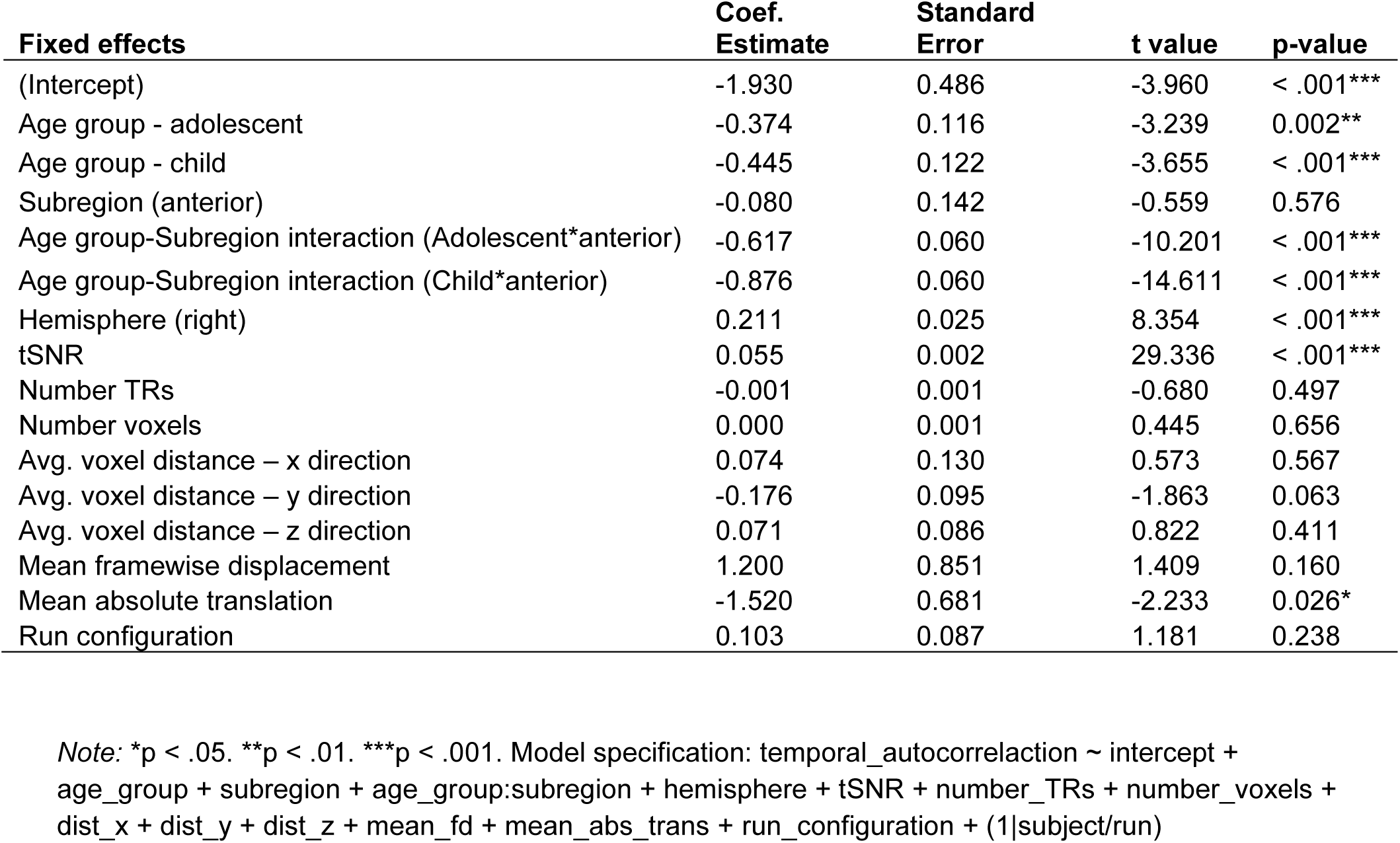
Differences in temporal autocorrelation by subregion and categorical age group in navigation task.

| **Fixed effects** | **Coef.**<br>**Estimate** | **Standard**<br>**Error** | **t value** | **p-value** |
| --- | --- | --- | --- | --- |
| (Intercept) | -1.930 | 0.486 | -3.960 | < .001*** |
| Age group - adolescent | -0.374 | 0.116 | -3.239 | 0.002** |
| Age group - child | -0.445 | 0.122 | -3.655 | < .001*** |
| Subregion (anterior) | -0.080 | 0.142 | -0.559 | 0.576 |
| Age group-Subregion interaction (Adolescent*anterior) | -0.617 | 0.060 | -10.201 | < .001*** |
| Age group-Subregion interaction (Child*anterior) | -0.876 | 0.060 | -14.611 | < .001*** |
| Hemisphere (right) | 0.211 | 0.025 | 8.354 | < .001*** |
| tSNR | 0.055 | 0.002 | 29.336 | < .001*** |
| Number TRs | -0.001 | 0.001 | -0.680 | 0.497 |
| Number voxels | 0.000 | 0.001 | 0.445 | 0.656 |
| Avg. voxel distance – x direction | 0.074 | 0.130 | 0.573 | 0.567 |
| Avg. voxel distance – y direction | -0.176 | 0.095 | -1.863 | 0.063 |
| Avg. voxel distance – z direction | 0.071 | 0.086 | 0.822 | 0.411 |
| Mean framewise displacement | 1.200 | 0.851 | 1.409 | 0.160 |
| Mean absolute translation | -1.520 | 0.681 | -2.233 | 0.026* |
| Run configuration | 0.103 | 0.087 | 1.181 | 0.238 |

*Note:* \*p < .05. \*\*p < .01. \*\*\*p < .001. Model specification: temporal\_autocorrelation ~ intercept + age\_group + subregion + age\_group:subregion + hemisphere + tSNR + number\_TRs + number\_voxels + dist\_x + dist\_y + dist\_z + mean\_fd + mean\_abs\_trans + run\_configuration + (1|subject/run)

**Table S7.** Follow-up analysis of continuous age differences in temporal autocorrelation within developmental group during navigation task by subregion.

| **Effect** | **Coef.**<br>**Estimate** | **Standard**<br>**Error** | **t value** | **p-value** |
| --- | --- | --- | --- | --- |
| Age (years and months) - Anterior | 0.118 | 0.033 | 3.592 | < .001*** |
| Age (years and months) - Posterior | 0.039 | 0.033 | 1.186 | 0.240 |

*Note:* \*p < .05. \*\*p < .01. \*\*\*p < .001. Slopes extracted directly from aggregate model (Table S6) using sim\_slopes function of interactions package in R.

**Table S8.** Temporal autocorrelation in parahippocampal cortex in task-based navigation sample.

| Fixed effects | Coef.<br>Estimate | Standard<br>Error | t value | p-value |
| --- | --- | --- | --- | --- |
| (Intercept) | -5.340 | 0.650 | -8.207 | < .001*** |
| Age group - adolescent | -0.068 | 0.115 | -0.588 | 0.558 |
| Age group - child | -0.187 | 0.120 | -1.557 | 0.123 |
| Hemisphere (right) | -0.135 | 0.037 | -3.649 | < .001*** |
| tSNR | 0.053 | 0.002 | 27.942 | < .001*** |
| Number TRs | -0.001 | 0.001 | -0.708 | 0.480 |
| Number voxels | -0.002 | 0.001 | -1.154 | 0.249 |
| Avg. voxel distance – x direction | 1.220 | 0.211 | 5.793 | < .001*** |
| Avg. voxel distance – y direction | 0.454 | 0.144 | 3.143 | 0.002** |
| Avg. voxel distance – z direction | -0.126 | 0.268 | -0.468 | 0.640 |
| Mean framewise displacement | 0.806 | 0.570 | 1.414 | 0.158 |
| Mean absolute translation | 0.500 | 0.596 | 0.839 | 0.402 |
| Run configuration | 0.102 | 0.095 | 1.068 | 0.286 |

*Note:* \*p < .05. \*\*p < .01. \*\*\*p < .001. Model specification: temporal\_autocorrelation ~ intercept + age\_group + hemisphere + tSNR + number\_TRs + number\_voxels + dist\_x + dist\_y + dist\_z + mean\_fd + mean\_abs\_trans + run\_configuration + (1|subject/run)

**Table S9.** Temporal autocorrelation during navigation task by age group while controlling for participant sex and general cognitive ability.

| Fixed effects | Coef.<br>Estimate | Standard<br>Error | t value | p-value |
| --- | --- | --- | --- | --- |
| (Intercept) | -1.890 | 0.631 | -3.001 | 0.003** |
| Vocabulary (WASI) | 0.008 | 0.006 | 1.400 | 0.166 |
| Sex (male) | -0.038 | 0.096 | -0.393 | 0.695 |
| Age group - adolescent | -0.447 | 0.127 | -3.514 | < .001*** |
| Age group - child | -0.551 | 0.136 | -4.053 | < .001*** |
| Subregion (anterior) | -0.348 | 0.153 | -2.269 | 0.023* |
| Age group-Subregion interaction (Adolescent*anterior) | -0.523 | 0.063 | -8.279 | < .001*** |
| Age group-Subregion interaction (Child*anterior) | -0.800 | 0.064 | -12.562 | < .001*** |
| Hemisphere (right) | 0.193 | 0.026 | 7.385 | < .001*** |
| tSNR | 0.056 | 0.002 | 28.755 | < .001*** |
| Number TRs | 0.000 | 0.001 | -0.359 | 0.720 |
| Number voxels | 0.001 | 0.001 | 1.447 | 0.148 |
| Avg. voxel distance – x direction | -0.121 | 0.138 | -0.875 | 0.382 |
| Avg. voxel distance – y direction | -0.325 | 0.100 | -3.244 | 0.001** |
| Avg. voxel distance – z direction | 0.118 | 0.089 | 1.328 | 0.184 |
| Mean framewise displacement | 1.610 | 0.925 | 1.742 | 0.082 |
| Mean absolute translation | -1.970 | 0.740 | -2.668 | 0.008** |
| Run configuration | 0.092 | 0.099 | 0.933 | 0.351 |

*Note:* \*p < .05. \*\*p < .01. \*\*\*p < .001. Model specification: temporal\_autocorrelation ~ intercept + WASI + sex + age\_group + subregion + age\_group:subregion + hemisphere + tSNR + number\_TRs + number\_voxels + dist\_x + dist\_y + dist\_z + mean\_fd + mean\_abs\_trans + run\_configuration + (1|subject/run)

**Table S10.**
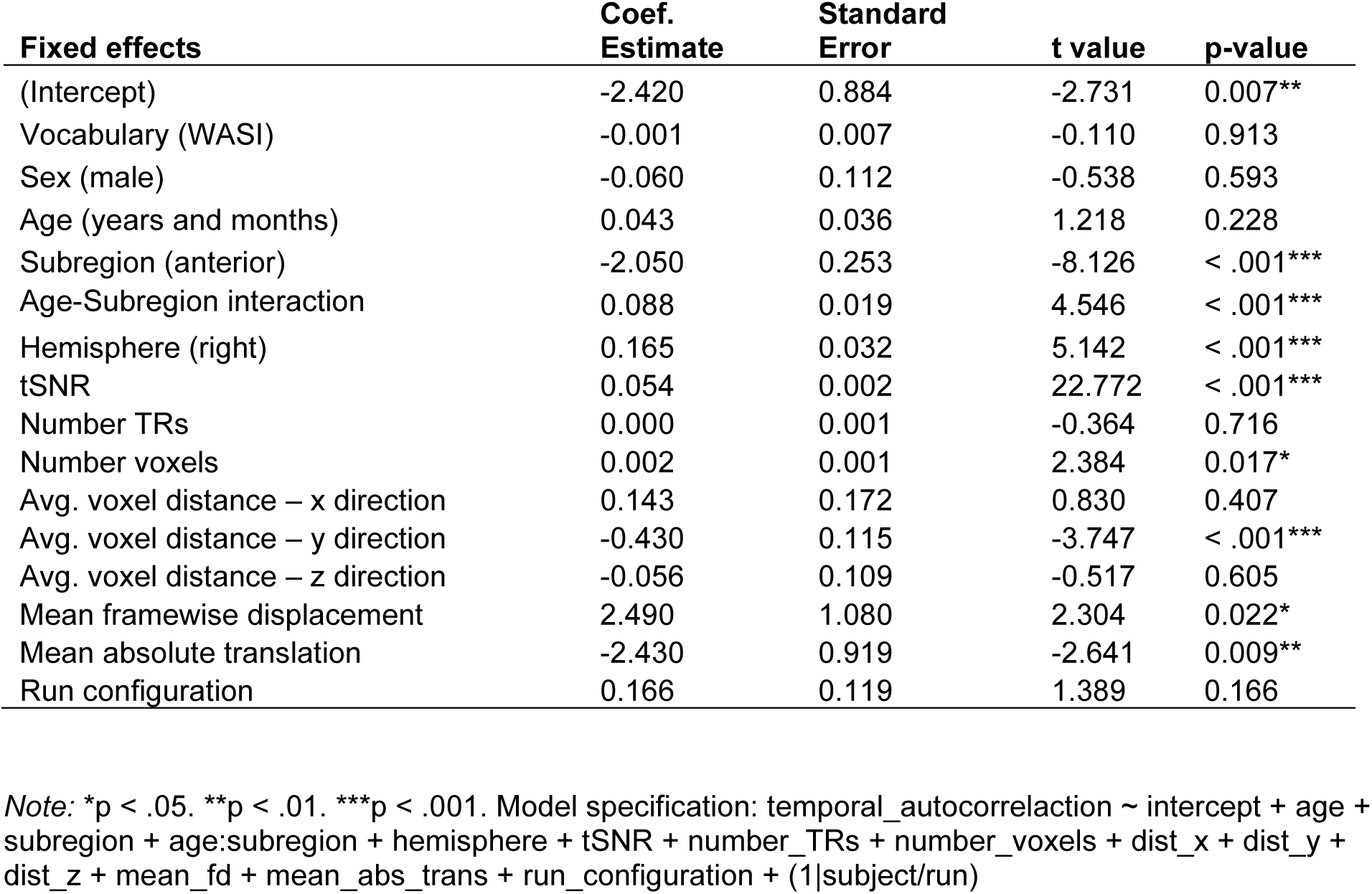
Temporal autocorrelation during navigation task by continuous age within developmental group while controlling for participant sex and general cognitive ability.

**Table S11.** Null effect of three-way interaction between scaled accuracy, age group, subregion.

| **Fixed effects** | **Coef.**<br>**Estimate** | **Standard**<br>**Error** | **t value** | **p-value** |
| --- | --- | --- | --- | --- |
| (Intercept) | -4.000 | 0.920 | -4.341 | < .001*** |
| Subregion (anterior) | -0.019 | 0.442 | -0.042 | 0.966 |
| Scaled accuracy | 0.028 | 0.010 | 2.710 | 0.008** |
| Age group - adolescent | 1.510 | 0.787 | 1.919 | 0.058 |
| Age group - child | 1.810 | 0.801 | 2.255 | 0.027* |
| Age group-Subregion interaction (Adolescent*anterior) | -0.924 | 0.436 | -2.120 | 0.034* |
| Age group-Subregion interaction (Child*anterior) | -1.400 | 0.440 | -3.181 | 0.002** |
| Behavior-Subregion interaction (Scaled accuracy *anterior) | -0.001 | 0.006 | -0.243 | 0.808 |
| Behavior-Age group interaction (Scaled accuracy *adolescent) | -0.025 | 0.011 | -2.356 | 0.021* |
| Behavior-Age group interaction (Scaled accuracy *child) | -0.033 | 0.011 | -2.848 | 0.006** |
| Behavior-Subregion-Age group interaction (Scaled accuracy *anterior *adolescent) | 0.005 | 0.006 | 0.816 | 0.415 |
| Behavior-Subregion-Age group interaction (Scaled accuracy *anterior *child) | 0.011 | 0.006 | 1.707 | 0.088 |
| Hemisphere (right) | 0.206 | 0.025 | 8.208 | < .001*** |
| tSNR | 0.055 | 0.002 | 29.464 | < .001*** |
| Number TRs | -0.001 | 0.001 | -0.632 | 0.528 |
| Number voxels | 0.000 | 0.001 | 0.456 | 0.649 |
| Avg. voxel distance – x direction | 0.111 | 0.131 | 0.851 | 0.395 |
| Avg. voxel distance – y direction | -0.216 | 0.095 | -2.276 | 0.023* |
| Avg. voxel distance – z direction | 0.082 | 0.088 | 0.940 | 0.348 |
| Mean framewise displacement | 1.440 | 0.848 | 1.699 | 0.090 |
| Mean absolute translation | -1.490 | 0.679 | -2.189 | 0.029* |
| Run configuration | 0.102 | 0.087 | 1.169 | 0.243 |

*Note:* \*p < .05. \*\*p < .01. \*\*\*p < .001. Model specification: temporal\_autocorrelation ~ intercept + subregion + scaled\_accuracy + age\_group + age\_group:subregion + scaled\_accuracy:subregion + scaled\_accuracy:age\_group + scaled\_accuracy:age\_group:subregion + hemisphere + tSNR + number\_TRs + number\_voxels + dist\_x + dist\_y + dist\_z + mean\_fd + mean\_abs\_trans + run\_configuration + (1|subject/run)

**Table S12.** Scaled accuracy and temporal autocorrelation during navigation task (reduced model)

| **Fixed effects** | **Coef.**<br>**Estimate** | **Standard**<br>**Error** | **t value** | **p-value** |
| --- | --- | --- | --- | --- |
| (Intercept) | -3.770 | 0.891 | -4.234 | < .001*** |
| Subregion (anterior) | -0.458 | 0.186 | -2.465 | 0.014* |
| Scaled accuracy | 0.025 | 0.010 | 2.487 | 0.015* |
| Age group - adolescent | 1.310 | 0.758 | 1.733 | 0.087 |
| Age group - child | 1.460 | 0.772 | 1.897 | 0.062 |
| Age group-Subregion interaction (Adolescent*anterior) | -0.544 | 0.065 | -8.385 | < .001*** |
| Age group-Subregion interaction (Child*anterior) | -0.734 | 0.075 | -9.760 | < .001*** |
| Behavior-Subregion interaction (Scaled accuracy *anterior) | 0.005 | 0.002 | 3.160 | 0.002** |
| Behavior-Age group interaction (Scaled accuracy *adolescent) | -0.023 | 0.010 | -2.208 | 0.030* |
| Behavior-Age group interaction (Scaled accuracy *child) | -0.027 | 0.011 | -2.462 | 0.016* |
| Hemisphere (right) | 0.208 | 0.025 | 8.288 | < .001*** |
| tSNR | 0.055 | 0.002 | 29.430 | < .001*** |
| Number TRs | -0.001 | 0.001 | -0.627 | 0.531 |
| Number voxels | 0.000 | 0.001 | 0.382 | 0.703 |
| Avg. voxel distance – x direction | 0.094 | 0.130 | 0.728 | 0.467 |
| Avg. voxel distance – y direction | -0.203 | 0.095 | -2.146 | 0.032* |
| Avg. voxel distance – z direction | 0.096 | 0.087 | 1.100 | 0.272 |
| Mean framewise displacement | 1.440 | 0.848 | 1.700 | 0.090 |
| Mean absolute translation | -1.490 | 0.679 | -2.190 | 0.029* |
| Run configuration | 0.102 | 0.087 | 1.168 | 0.244 |

*Note:* \*p < .05. \*\*p < .01. \*\*\*p < .001. Model specification: temporal\_autocorrelation ~ intercept + subregion + navigation\_accuracy + age\_group + age\_group:subregion + navigation\_accuracy:subregion + navigation\_accuracy:age\_group + hemisphere + tSNR + number\_TRs + number\_voxels + dist\_x + dist\_y + dist\_z + mean\_fd + mean\_abs\_trans + run\_configuration + (1|subject/run)

**Table S13.** Raw (non-scaled) distance error and temporal autocorrelation during navigation task.

| **Fixed effects** | **Coef.**<br>**Estimate** | **Standard**<br>**Error** | **t value** | **p-value** |
| --- | --- | --- | --- | --- |
| (Intercept) | -1.330 | 0.539 | -2.460 | 0.014* |
| Subregion (anterior) | 0.028 | 0.145 | 0.195 | 0.845 |
| Distance error (raw) | -0.080 | 0.032 | -2.470 | 0.016* |
| Age group - adolescent | -0.953 | 0.303 | -3.141 | 0.002** |
| Age group - child | -1.220 | 0.387 | -3.164 | 0.002** |
| Age group - Subregion interaction (Adolescent*anterior) | -0.538 | 0.065 | -8.281 | < .001*** |
| Age group - Subregion interaction (Child*anterior) | -0.723 | 0.075 | -9.599 | < .001*** |
| Behavior - Subregion interaction (Distance error*anterior) | -0.016 | 0.005 | -3.366 | 0.001** |
| Behavior - Age group interaction (Distance error*adolescent) | 0.074 | 0.033 | 2.200 | 0.031* |
| Behavior - Age group interaction (Distance error*child) | 0.086 | 0.036 | 2.420 | 0.018* |
| Hemisphere (right) | 0.208 | 0.025 | 8.298 | < .001*** |
| tSNR | 0.055 | 0.002 | 29.435 | < .001*** |
| Number TRs | -0.001 | 0.001 | -0.600 | 0.549 |
| Number voxels | 0.000 | 0.001 | 0.347 | 0.728 |
| Avg. voxel distance – x direction | 0.095 | 0.129 | 0.731 | 0.465 |
| Avg. voxel distance – y direction | -0.203 | 0.095 | -2.144 | 0.032* |
| Avg. voxel distance – z direction | 0.099 | 0.087 | 1.133 | 0.258 |
| Mean framewise displacement | 1.460 | 0.848 | 1.719 | 0.086 |
| Mean absolute translation | -1.490 | 0.679 | -2.193 | 0.029* |
| Run configuration | 0.101 | 0.087 | 1.161 | 0.246 |

*Note:* \*p < .05. \*\*p < .01. \*\*\*p < .001. Model specification: temporal\_autocorrelation ~ intercept + subregion + distance\_error + age\_group + age\_group:subregion + distance\_error:subregion + distance\_error:age\_group + hemisphere + tSNR + number\_TRs + number\_voxels + dist\_x + dist\_y + dist\_z + mean\_fd + mean\_abs\_trans + run\_configuration + (1|subject/run)

**Table S14.** Follow-up analysis of age group differences in relation between scaled accuracy and temporal autocorrelation by subregion.

| **Effect** | **Coef. Estimate** | **Standard Error** | **t value** | **p-value** |
| --- | --- | --- | --- | --- |
| Adult - Anterior | 0.029 | 0.010 | 2.974 | 0.004** |
| Adult - Posterior | 0.025 | 0.010 | 2.487 | 0.015* |
| Adolescent - Anterior | 0.007 | 0.003 | 2.142 | 0.035* |
| Adolescent - Posterior | 0.002 | 0.003 | 0.604 | 0.548 |
| Child - Anterior | 0.002 | 0.005 | 0.466 | 0.643 |
| Child - Posterior | -0.003 | 0.005 | -0.503 | 0.616 |

*Note:* \* $p < .05$ . \*\* $p < .01$ . \*\*\* $p < .001$ . Slopes extracted directly from aggregate model (Table S12) using `sim_slopes` function of `interactions` package in R.

**Table S15.**
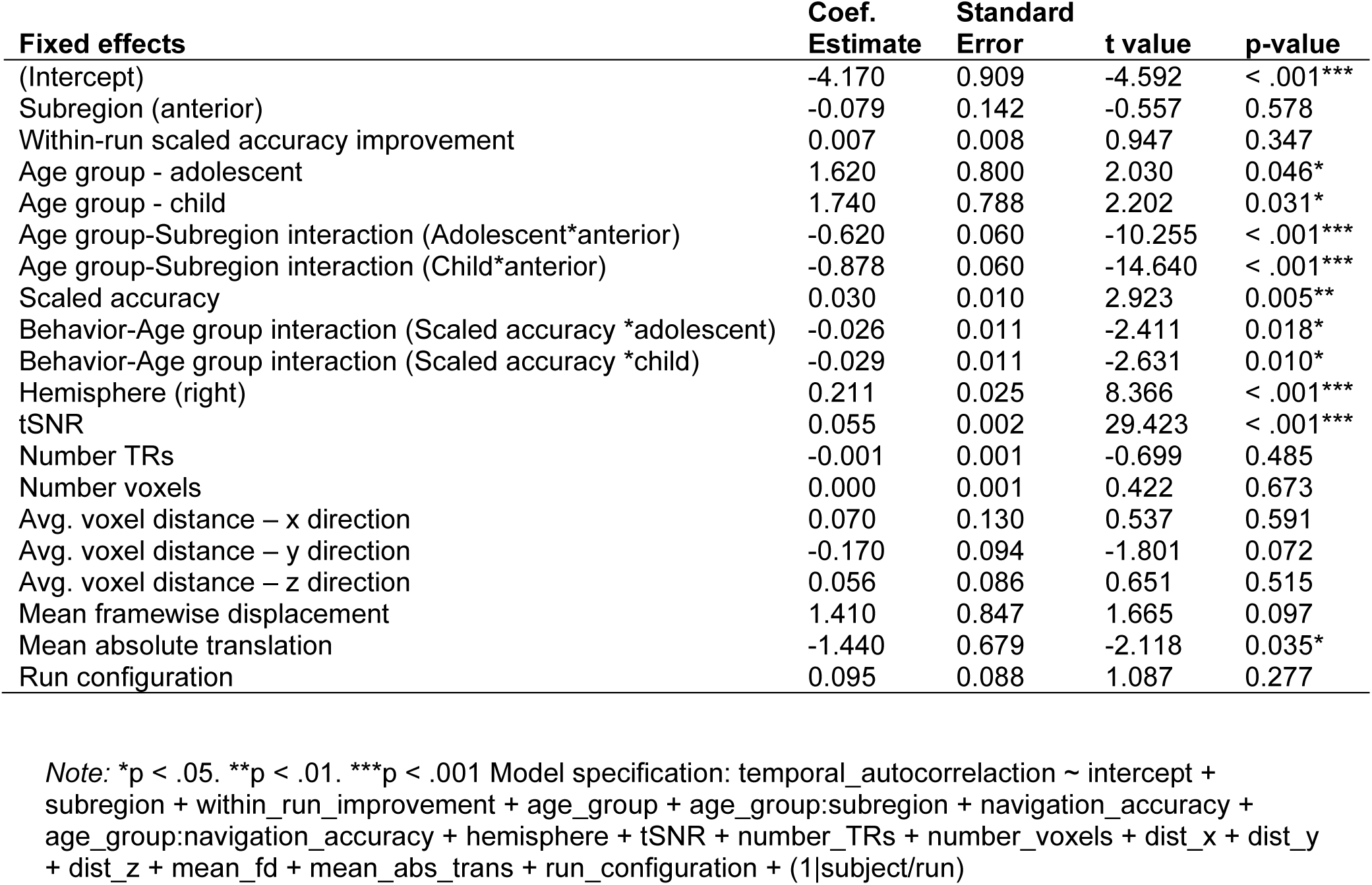
Within-run scaled accuracy improvement and temporal autocorrelation during navigation task.

| **Fixed effects** | **Coef.**<br>**Estimate** | **Standard**<br>**Error** | **t value** | **p-value** |
| --- | --- | --- | --- | --- |
| (Intercept) | -4.170 | 0.909 | -4.592 | < .001*** |
| Subregion (anterior) | -0.079 | 0.142 | -0.557 | 0.578 |
| Within-run scaled accuracy improvement | 0.007 | 0.008 | 0.947 | 0.347 |
| Age group - adolescent | 1.620 | 0.800 | 2.030 | 0.046* |
| Age group - child | 1.740 | 0.788 | 2.202 | 0.031* |
| Age group-Subregion interaction (Adolescent*anterior) | -0.620 | 0.060 | -10.255 | < .001*** |
| Age group-Subregion interaction (Child*anterior) | -0.878 | 0.060 | -14.640 | < .001*** |
| Scaled accuracy | 0.030 | 0.010 | 2.923 | 0.005** |
| Behavior-Age group interaction (Scaled accuracy *adolescent) | -0.026 | 0.011 | -2.411 | 0.018* |
| Behavior-Age group interaction (Scaled accuracy *child) | -0.029 | 0.011 | -2.631 | 0.010* |
| Hemisphere (right) | 0.211 | 0.025 | 8.366 | < .001*** |
| tSNR | 0.055 | 0.002 | 29.423 | < .001*** |
| Number TRs | -0.001 | 0.001 | -0.699 | 0.485 |
| Number voxels | 0.000 | 0.001 | 0.422 | 0.673 |
| Avg. voxel distance – x direction | 0.070 | 0.130 | 0.537 | 0.591 |
| Avg. voxel distance – y direction | -0.170 | 0.094 | -1.801 | 0.072 |
| Avg. voxel distance – z direction | 0.056 | 0.086 | 0.651 | 0.515 |
| Mean framewise displacement | 1.410 | 0.847 | 1.665 | 0.097 |
| Mean absolute translation | -1.440 | 0.679 | -2.118 | 0.035* |
| Run configuration | 0.095 | 0.088 | 1.087 | 0.277 |

*Note:* \*p < .05. \*\*p < .01. \*\*\*p < .001 Model specification: temporal\_autocorrelation ~ intercept + subregion + within\_run\_improvement + age\_group + age\_group:subregion + navigation\_accuracy + age\_group:navigation\_accuracy + hemisphere + tSNR + number\_TRs + number\_voxels + dist\_x + dist\_y + dist\_z + mean\_fd + mean\_abs\_trans + run\_configuration + (1|subject/run)

**Table S16.** Excess path and temporal autocorrelation during navigation task.

| **Fixed effects** | **Coef.**<br>**Estimate** | **Standard**<br>**Error** | **t value** | **p-value** |
| --- | --- | --- | --- | --- |
| (Intercept) | 2.710 | 7.740 | 0.350 | 0.727 |
| Subregion (anterior) | 0.772 | 0.210 | 3.682 | <.001*** |
| Excess path | -0.016 | 0.019 | -0.854 | 0.395 |
| Age group - adolescent | -0.152 | 0.388 | -0.393 | 0.696 |
| Age group - child | -0.736 | 0.409 | -1.800 | 0.075 |
| Age group-Subregion interaction (Adolescent*anterior) | -1.630 | 0.212 | -7.702 | <.001*** |
| Age group-Subregion interaction (Child*anterior) | -1.570 | 0.220 | -7.130 | <.001*** |
| Behavior-Subregion interaction (Excess path *anterior) | -0.057 | 0.010 | -5.796 | <.001*** |
| Behavior-Age group interaction (Excess path *adolescent) | -0.002 | 0.020 | -0.087 | 0.931 |
| Behavior-Age group interaction (Excess path *child) | 0.016 | 0.020 | 0.829 | 0.410 |
| Behavior-Subregion-Age group interaction (Excess path *anterior *adolescent) | 0.059 | 0.011 | 5.459 | <.001*** |
| Behavior-Subregion-Age group interaction (Excess path *anterior *child) | 0.050 | 0.011 | 4.661 | <.001*** |
| Scaled accuracy | -0.037 | 0.084 | -0.445 | 0.658 |
| Corrective Feedback | -0.122 | 0.270 | -0.450 | 0.654 |
| Average Euclidean path | -0.017 | 0.033 | -0.518 | 0.606 |
| Hemisphere (right) | 0.209 | 0.025 | 8.248 | <.001*** |
| tSNR | 0.056 | 0.002 | 29.783 | <.001*** |
| Number TRs | 0.000 | 0.001 | 0.414 | 0.679 |
| Number voxels | 0.000 | 0.001 | 0.473 | 0.636 |
| Avg. voxel distance – x direction | 0.055 | 0.131 | 0.416 | 0.677 |
| Avg. voxel distance – y direction | -0.199 | 0.095 | -2.094 | 0.036* |
| Avg. voxel distance – z direction | 0.005 | 0.088 | 0.058 | 0.954 |
| Mean framewise displacement | 1.480 | 0.845 | 1.751 | 0.081 |
| Mean absolute translation | -1.550 | 0.674 | -2.299 | 0.022* |
| Run configuration | 0.093 | 0.086 | 1.082 | 0.280 |

*Note:* \*p < .05. \*\*p < .01. \*\*\*p < .001. Model specification: temporal\_autocorrelation ~ intercept + subregion + excess\_path + age\_group + age\_group:subregion + excess\_path:subregion + excess\_path:age\_group + excess\_path:subregion:age\_group + navigation\_accuracy + corrective\_feedback + Euclidean\_path hemisphere + tSNR + number\_TRs + number\_voxels + dist\_x + dist\_y + dist\_z + mean\_fd + mean\_abs\_trans + run\_configuration + (1|subject/run)

**Table S17.** Follow-up analysis of age group differences in relation between excess path and temporal autocorrelation by subregion.

| **Effect** | **Coef.**<br>**Estimate** | **Standard**<br>**Error** | **t value** | **p-value** |
| --- | --- | --- | --- | --- |
| Adult - Anterior | -0.072 | 0.018 | -3.918 | <.001*** |
| Adult - Posterior | -0.016 | 0.019 | -0.854 | 0.395 |
| Adolescent - Anterior | -0.015 | 0.010 | -1.443 | 0.153 |
| Adolescent - Posterior | -0.018 | 0.010 | -1.725 | 0.088 |
| Child - Anterior | -0.006 | 0.009 | -0.700 | 0.486 |
| Child - Posterior | 0.001 | 0.009 | 0.067 | 0.947 |

*Note:* \*p < .05. \*\*p < .01. \*\*\*p < .001. Slopes extracted directly from aggregate model (Table S16) using sim\_slopes function of interactions package in R.

## Notes

https://osf.io/g7qne

### Competing Interest Statement

The authors have declared no competing interest.

### Summary of Updates

Manuscript updated in response to reviewer comments.

